# Spatial and temporal muscle synergies provide a dual characterization of low-dimensional and intermittent control of upper-limb movements

**DOI:** 10.1101/2022.07.11.499519

**Authors:** Cristina Brambilla, Manfredo Atzori, Henning Müller, Andrea d’Avella, Alessandro Scano

## Abstract

Muscle synergy analysis is commonly used for investigating the neurophysiological mechanisms that the central nervous system employs to control muscle activations. In the last two decades, several models have been developed to decompose EMG signals into spatial, temporal or spatiotemporal synergies. However, the presence of different approaches complicates the comparison and interpretation of results. Spatial synergies represent invariant activation weights in muscle groups modulated with variant temporal coefficients, while temporal synergies are based on invariant temporal profiles that coordinate variant muscle weights. While non-negative matrix factorization (NMF) allows to extract both spatial and temporal synergies, temporal synergies and the comparison between the two approaches have been barely investigated and so far no study targeted a large set of multi-joint upper limb movements. Here we present several analyses that highlight the duality of spatial and temporal synergies as a characterization of low-dimensional and intermittent motor coordination in the upper limb, allowing high flexibility and dexterity. First, spatial and temporal synergies were extracted from two datasets representing a comprehensive mapping of proximal (REACH PLUS) and distal (NINAPRO) upper limb movements, focusing on their differences in reconstruction accuracy and inter-individual variability. For both models, we extracted synergies achieving a given level of the goodness of reconstruction (*R*^*2*^), and we compared the similarity of the invariant components across participants. The two models provide a compact characterization of motor coordination at spatial or temporal level, respectively. However, a lower number of temporal synergies are needed to achieve the same *R*^*2*^ with a higher inter-subject similarity. Spatial and temporal synergies may thus capture different levels of motor control. Second, we showed the existence of both spatial and temporal structure in the EMG data, extracting spatial and temporal synergies from a surrogate dataset in which the phases were shuffled preserving the same frequency content of the original data. Last, a detailed characterization of the structure of the temporal synergies suggested that they can be related to an intermittent control of the movement. These results may be useful to improve muscle synergy analysis in several fields such as rehabilitation, prosthesis control and motor control studies.

## 1. Introduction

The study of motor control focuses on how the central nervous system (CNS) executes and coordinates complex movements involving several muscles. Human motor control relies on a combination of a limited number of spatial and/or temporal patterns or modules (Bizzi et al., 2008), to simplify the planning and the production of movement. These patterns of muscular activations are often referred to as muscle synergies. In the last two decades, many studies have exploited different approaches based on muscle synergies for the analysis of human motor control and different models have been developed to decompose the EMG signals into spatial, temporal or spatiotemporal organizations. Existing models are spatial or synchronous synergies (Cheung et al., 2005; Ting & Macpherson, 2005; Tresch et al., 1999), invariant temporal components or temporal synergies (Ivanenko et al., 2004; 2005), spatiotemporal or time-varying synergies (D’Avella et al., 2003, 2006) and space-by-time synergistic models (Delis et al., 2014, 2015).

Spatial and temporal synergies are based on simple invariant modules: in spatial synergies, invariant muscle weights are modulated by variant temporal coefficients; in temporal synergies, temporal invariant synergies modulate variant muscle weights. Typically, muscle patterns observed in different conditions of a motor task are captured by the variable combination of invariant spatial or temporal synergies. The spatiotemporal model on the other hand captures the variability in the muscle patterns as due to the amplitude modulation and temporal delay of invariant collections of potentially asynchronous muscle activation waveforms. The space-by-time model proposes invariant spatial and temporal synergies combined into invariant synchronous spatiotemporal patterns that are recruited by variant coefficients. In summary, all synergy models assume that a few invariant modules are recruited with some spatially or temporally variant coefficients to account for the variability observed in the muscle patterns.

The modules, i.e., the muscle synergies, however, may also vary. Muscle synergy variability can be affected by many factors, such as the synergy model used, and the algorithm chosen for the extraction. Spatial and temporal synergies can be extracted with NMF (Lee & Seung, 1999). Interestingly, the standard NMF is mostly used to extract spatial synergies rather than temporal synergies. This is probably due to the fact that original work on muscle synergies focused on the spinal cord and demonstrated a spatial organization of muscle patterns (Cheung et al., 2005; Tresch et al., 1999). Another factor that influences muscle synergy variability is whether EMG data are averaged over repetitions of the same task or concatenated. For the spatial model, considering *M* muscles, *K* tasks sampled with *T* samples each, and *R* repetitions of each group of tasks, and *S* extracted synergies, the EMG signals are grouped in a data matrix with *M* rows and *K·T* columns, if the task repetitions are averaged; on the contrary, if task repetitions are concatenated (Oliveira et al., 2014), EMG signals are grouped in a data matrix with *M* rows and *K·T·R* columns. Extracted synergies are thus extracted as a matrix with *M* rows and *S* columns of invariant spatial synergies modulated by variant temporal coefficients typical of each task. While this model is by far the most frequently employed in the literature, the dual module (time invariant temporal synergies) may also be used. For the temporal model, the EMG signals are grouped in a data matrix with *T* rows and *K·M* columns, if the task repetitions are averaged; on the contrary, if task repetitions are concatenated, EMG signals are grouped in a data matrix with *T* rows and *K·M·R* columns. Extracted synergies are thus organized in a matrix with *T* rows and *S* columns of invariant temporal synergies modulated by variant muscle weights typical of each task. A schematic illustration of how the matrix is arranged for each case is shown in Figure 1.

**Figure 1.**
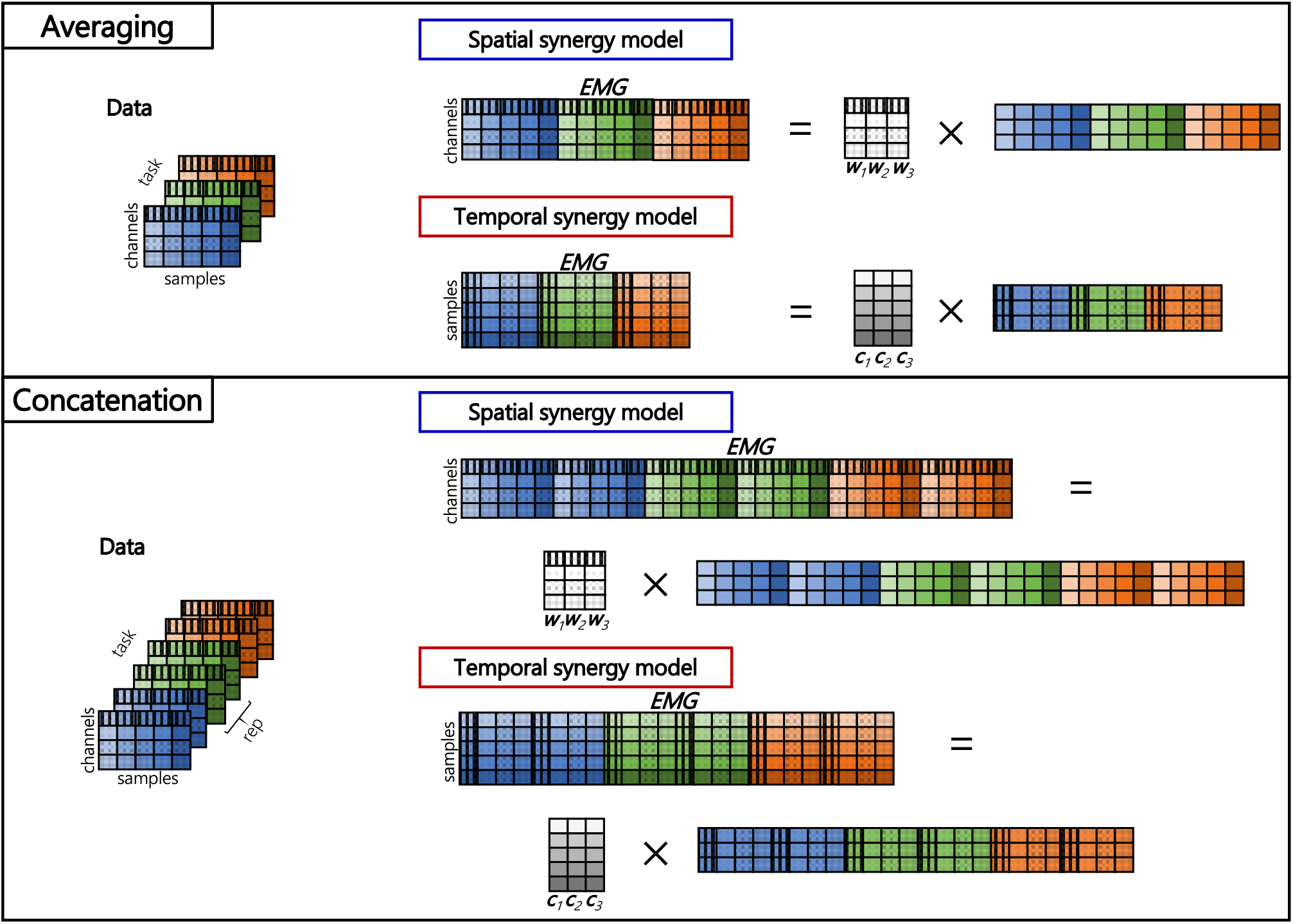
Schematic representation of data matrix arrangement and decomposition for both spatial and temporal synergy models. In the upper panel, task repetitions are averaged, while in the lower panel the repetitions are concatenated. Data are collected in matrices in which the task is represented by the color (blue, green, orange), the channels (4 in the example) are represented by different patterns and the time samples (5) are represented by different color saturation levels.

The temporal model is based on the hypothesis that activation profiles - rather than spatial weights - are invariant. The extraction of this type of synergies is usually employed in periodic tasks, like locomotion (Ivanenko et al., 2004, 2005) and cycling (Torricelli et al., 2020), where an invariant temporal structure is naturally found. However, the typical, highly repeatable bell-shaped velocity profiles found in human upper limb discrete movement (Flash & Hogan, 1985) also suggest the existence of temporal modules, in addition to spatial modules, whose amplitude modulation explains the observed directional tuning of muscle activations (Borzelli et al., 2013; Mira et al., 2021; Scano et al., 2019). We argue that spatial synergies may characterize motor coordination well at the spinal (Takei et al., 2017) and the corticospinal (Cheung et al., 2009, 2012) levels, while temporal and spatiotemporal synergies may be employed at the cortical (Overduin et al., 2015) and the cerebellar (Berger et al., 2020) levels for coordination even at higher levels of the neuro-motor hierarchy (Wolpert & Kawato, 1998).

The presence of multiple definitions of muscle synergies in the literature complicates the comparison and the interpretation of the results obtained from different studies. Few studies are available that compare the spatial and the temporal models and show what their differences imply in the interpretation of the data. Delis and collaborators (2014) compared the spatial and temporal models with the space-by-time model and found that the space-by-time model, while compatible with the two models, provides a more parsimonious representation of muscle activation patterns. Chiovetto and colleagues (2013) compared temporal, spatial and spatiotemporal models by extracting muscle synergies from single joint movements, including only two muscles. They showed that all the three models lead to interpretable synergies that encode specific motor features: in particular, spatial synergies describe the coordinated activation of a group of muscles, while the temporal ones reveal the different phases of the movement. Safavynia and Ting (2012) used both spatial and temporal synergies to analyze postural control under perturbation and found that spatial synergies reconstruct the EMG signals better and are more interpretable. Russo and collaborators (2014) employed the spatial, temporal and spatiotemporal synergy models to investigate and compare the dimensionality of joint torques to the one of muscle patterns. They found that joint torques have a lower dimensionality with respect to muscles and the temporal synergy model is more parsimonious than the spatial one. Spatial and temporal synergy models were also compared by Torricelli and collaborators (2020), who also used a surrogate dataset to evaluate the process of short-term adaptation in cycling tasks. Their results showed that neither spatial nor temporal model could describe the learning process adequately, while the temporal model shifted by an optimal delay could explain the changes in muscle coordination. Finally, Berger et al. (2020) studied the role of the cerebellum on the organization of movement employing spatial, temporal and spatiotemporal synergy models and found that spatiotemporal synergies could identify changes in muscular pattern as a specific effect of cerebellar damage.

Despite the high number of studies in the domain, there are still open challenges. First, while the temporal model has been employed less frequently than the spatial one, no study to date has investigated at what extent temporal synergies reflect a specific temporal structure of the muscle activation patterns rather than only their intrinsic smoothness. Second, while several studies compared spatial and temporal synergies, there are not studies in literature targeting a large set of multi-joint upper limb movements.

To address these issues, this paper aims at assessing the existence of significative spatial and temporal structure and at comparing the spatial and temporal models when using the same EMG datasets. Two comprehensive datasets were used that include many muscles and degrees of freedom of the upper limb. The first one includes proximal upper-limb muscles (Scano et al., 2019) and the other distal upper-limb muscles (Atzori et al., 2014). The first objective was achieved building a surrogate dataset from the original data, randomly shuffling the phases of the Fourier Transform of the signal and preserving the frequency content (Torricelli et al., 2020). The spatial and the temporal synergy models were employed on both the original dataset and the surrogate one and the goodness of the reconstruction was compared in order to find significative differences between the synergies extracted from original and surrogate datasets. After validating the synergy models, spatial and temporal synergies were extracted with NMF from both original datasets and a comprehensive comparison was performed. The reconstruction capability of the two models suggests a different organization of the motor control between proximal and distal upper limbs. Differences in inter-subject variability of the invariant components between the two models and the two datasets were observed. Furthermore, we provide a detailed characterization of the temporal features of the temporal synergies, suggesting that they can be related to an intermittent control scheme of the movement that allows high flexibility and dexterity.

## 2. Materials & Methods

An overview of the study is shown in Figure 2.

**Figure 2.**
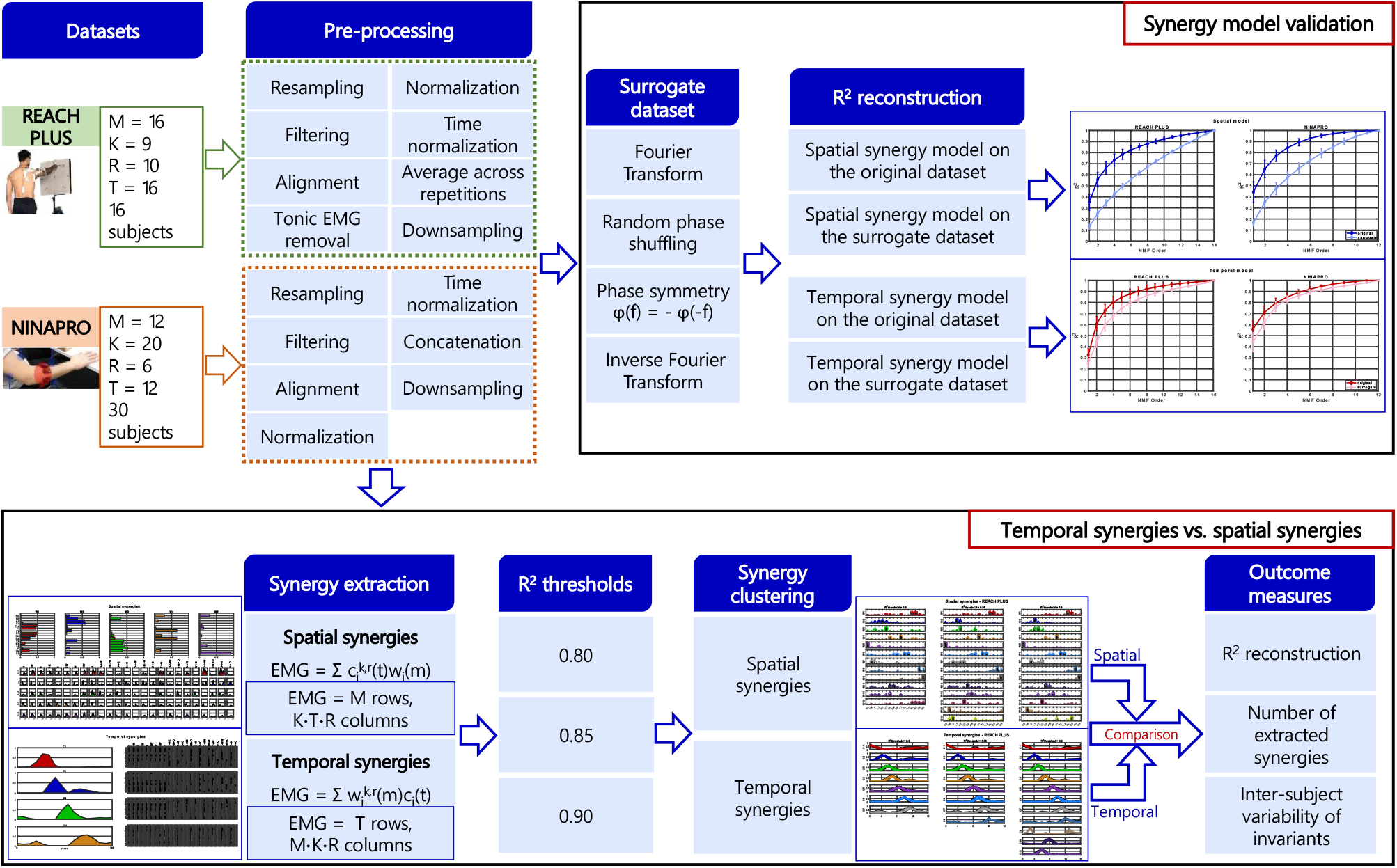
Schematic of the workflow for the analysis of both REACH PLUS and NINAPRO datasets. Each dataset is composed of M muscles, K tasks, R repetitions and T time samples. In the first row, the workflow for the validation of the synergy model is reported, while the comparison between temporal and spatial synergy model is shown in the second row.

### 2.1 Dataset description and preprocessing

Two datasets were selected for this analysis: the REACH PLUS dataset, representing the proximal upper-limb coordination in multi-directional movements (Scano et al., 2019) and the publicly available distal upper-limb dataset NINAPRO (Atzori et al., 2014). They both were already used for muscle synergy extraction with the spatial synergy model based on NMF (Pale et al., 2020; Scano et al., 2018).

The REACH PLUS dataset features 16 healthy participants performing frontal point to point movements towards 9 main cardinal directions and frontal exploration tasks from a central point towards 8 cardinal directions, and back to the central point; each group of movements was repeated ten times. The EMG signal was recorded from 16 muscles of the right upper limb. The data pre-processing was achieved with the methods already described in detail in a previous report (Scano et al., 2019). In summary, EMG data were resampled, filtered, aligned and the tonic component was removed with a linear ramp model. Normalization was performed on the maximum value of filtered EMG for each channel. The data from each phase were down sampled to 16 samples, so that spatial and temporal synergies were extracted from the same total number of samples.

The NINAPRO dataset features 30 selected healthy participants performing 20 hand grasps, extracted from hand repertoire of NINAPRO Dataset 2 (NINAPRO DB2). Each group of movements was repeated six times. Twelve channels were used for the recording of the EMG signal. The data pre-processing was achieved with the methods described in detail in a previous report (Scano et al., 2018): EMG data were resampled, filtered, aligned and finally normalized on the maximum value of filtered EMG for each channel. The data from each phase were down sampled to 12 samples, so that spatial and temporal synergies were extracted from the same total number of samples.

The two datasets were analyzed separately and the results were compared.

### 2.2 Surrogate dataset

A preliminary step for our analysis was to validate both the spatial and temporal models for synergy extraction, demonstrating that a significative spatial and temporal organization exists in the muscle activation patterns and, in particular, that the spatial synergies do not simply represent a feature related to the amplitude distribution of the signals and the temporal synergies a feature related to the smoothness of the signals. To do this, we constructed a surrogate dataset mimicking the amplitude distribution and smoothness of the original dataset (having the same Fourier components) but removing its specific spatiotemporal structure (by randomization of the phases of the Fourier components of each muscle), inspired from a previously adopted procedure (Torricelli et al., 2020). Following the approach of Faes et al. (2004), the surrogate datasets were constructed computing the Fourier Transform (FT) of the original time series; for each frequency component, we substituted the phases with random phases *ϕ* chosen in the interval [−*π, π*], while the modulus remained unchanged. Therefore, each complex amplitude obtained from FT was multiplied by *e*^*iϕ*^ and, in order to compute a real inverse FT, the phases were symmetrized to have *ϕ*(*f*) = −*ϕ*(−*f*) (Theiler et al., 1992). Finally, the inverse of the Fourier Transform was applied to return into the time domain, obtaining time series with the same frequency content of the original data but with random temporal structure. An example of the original and the surrogate dataset is shown in Figure 3. These datasets were given as input to the spatial and the temporal synergy extraction algorithm and the resulting reconstruction R^2^ and order of factorization for each R^2^ threshold were compared to those obtained from the original dataset.

**Figure 3.**
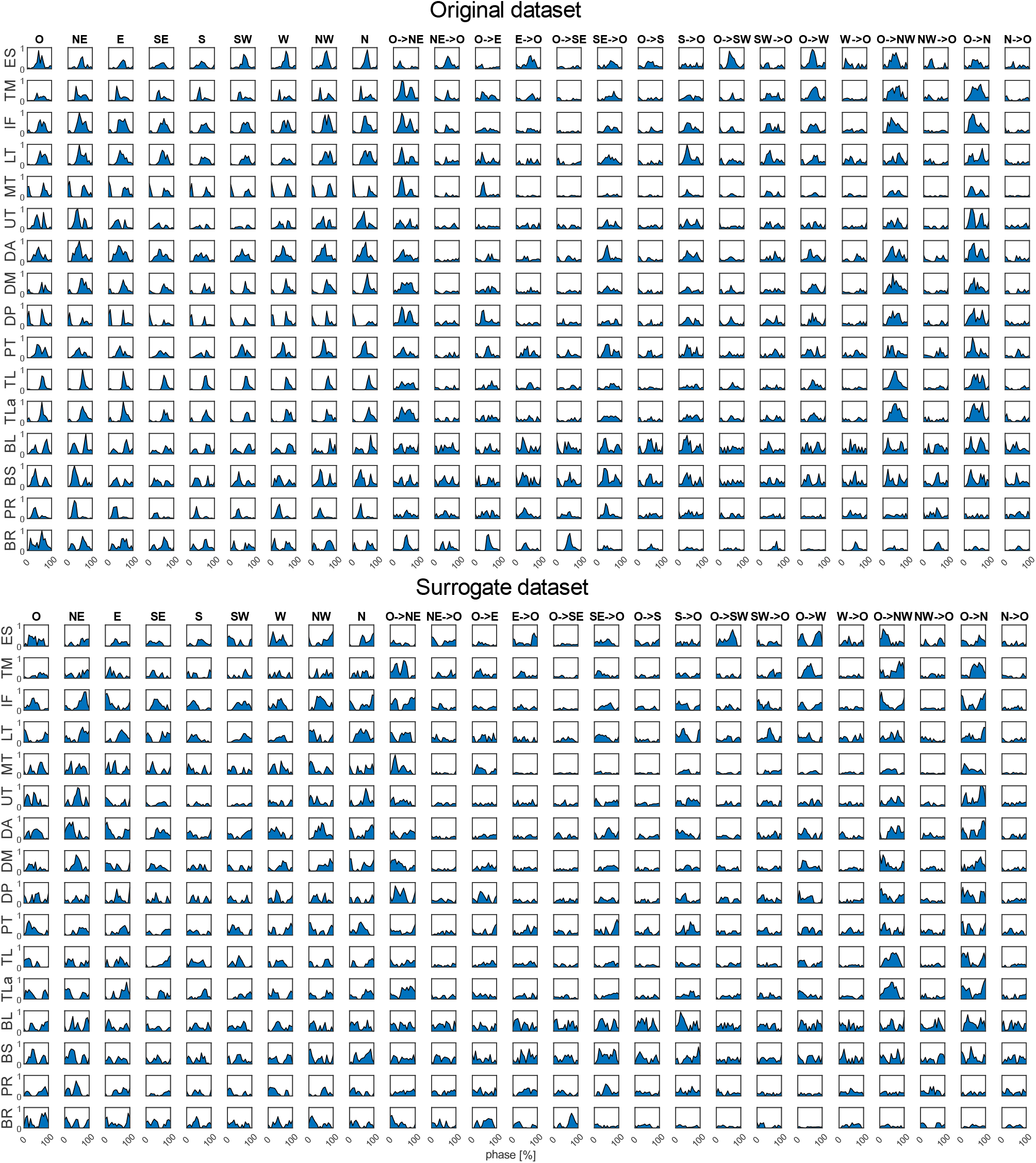
Example of the original (upper panel) and the surrogate (lower panel) dataset from one subject of the REACH PLUS dataset.

The bootstrapping procedure was performed for both datasets and repeated ten times for each participant: therefore, ten surrogate datasets were obtained and only the mean R^2^ over the ten repetitions for each participant was taken into consideration.

### 2.3 Synergy extraction

As in a previous study (Scano et al., 2019), data from the REACH PLUS dataset were averaged across repetitions and arranged differently for the extraction of spatial and temporal synergies. For extracting spatial synergies, considering *M* muscles, *K* tasks sampled with *T* samples each, the EMG signals were rearranged in a data matrix with *M* rows and *K* · *T* columns. In contrast, for extracting temporal synergies, the EMG data matrix had *T* rows and *K* · *M* columns.

Since there are 6 repetitions of each movement in the NINAPRO dataset and they had a longer duration, data were concatenated in order to preserve the trial by trial variability (Oliveira et al., 2014). Therefore, considering *M* muscles, *K* tasks sampled with *T* samples each, and *R* repetitions of each group of tasks, the EMG signals were arranged in a data matrix with *M* rows and *K* · *T* · *R* columns for extracting spatial synergies and in a data matrix with *T* rows and *K* · *R* · *M* columns for extracting temporal synergies.

For extracting both spatial and temporal synergies, we used a non-negative matrix factorization algorithm. The spatial model is the following:

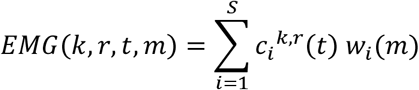

where ***w***_***i***_ are the time-invariant synergy vectors and ***c***_***i***_ the time-varying scalar activation coefficients for each synergy (*i* = 1…*S*), and *EMG*(*t*, k) the activity of muscle *m* at time *t* of task *k*.

For the REACH PLUS dataset, considering *K* = 25, *T* = 16, *M* = 16, input data was a 16 · 400 matrix. Spatial extraction would lead to *S* synergies (each a column vector with 16 components) and *S* · 25 time-varying coefficients (16 samples each). For NINAPRO dataset, input data was a 12 · 1440 matrix, considering *K* = 20, *T* = 12, *R* = 6, *M* = 12. Spatial extraction would lead to *S* synergies (each a column vector with 12 components) and *S* · 120 time-varying coefficients (12 samples each). Each spatial synergy was normalized by the Euclidean norm of that synergy and as a consequence the temporal coefficients were also normalized by the reciprocal of the norm.

The temporal model is:

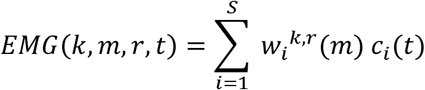

Where ***c***_***i***_ are the invariant temporal synergy vectors and ***w***_***i***_ the variant muscle weight vectors for each synergy dependent on task and repetition. Using the REACH PLUS dataset, input data was a 16 · 400 matrix and the extraction would lead to *S* temporal synergies (16 · *S* matrix) and *S* · 400 variant muscle weights (25 loads per spatial group). With NINAPRO dataset, input data was a 12 · 1440 matrix and the extraction would lead to *S* temporal synergies (12 · *S* matrix) and *S* · 1440 variant muscle weights (120 weights per spatial group).

The order of factorization *S*, given as input in the NMF algorithm, was increased from 1 to the number of EMG channels (16 for REACH PLUS and 12 for NINAPRO) for both spatial and temporal synergy extraction. For each *S*, the algorithm was applied 100 times with different random initializations in order to avoid local minima and the optimization run accounting for the higher variance of the signal was chosen as the representative of the order *S*.

As a measure of goodness of reconstruction, we used the *R*^2^ defined as 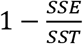 where *SSE* is the sum of the squared residuals and *SST* is the sum of the squared differences with the mean EMG vector (D’Avella et al., 2006). The order of factorization was chosen as the lowest one explaining a predefined threshold level of R^2^. We chose three threshold levels commonly adopted in literature (Pale et al., 2020) that were 0.80, 0.85 and 0.90.

### 2.4 Synergy clustering

To evaluate the variability of synergies from different participants, we grouped invariant spatial and temporal synergies across participants for each *R*^*2*^ level. Cluster analysis allows to reduce the dataset of extracted synergies to a limited number of groups, which compactly represent the repertoire of modules extracted among the participants. Since the order of factorization varied across the participants, we decided to group the extracted synergies using the k-means clustering algorithm (Steele et al., 2015).

A matrix containing the whole set of muscle synergies extracted from all the participants was given as input to the algorithm. The number of clusters was initially set equal to the maximum order of factorization obtained among the participants for the chosen threshold and was increased until all the synergies from the same participant were assigned to different clusters. The number of replicates was set to 200, namely the number of times the algorithm repeated the clustering with new initial cluster centroids estimates (chosen uniformly at random (Arthur & Vassilvitskii, 2007)) with the same number of clusters and give the results with the lowest sum of Euclidean distances of each point in the cluster to the centroid. The entire procedure was repeated 10 times and we considered as the best solution the one that gave the most parsimonious number of clusters. The algorithm gave the synergies grouped in each cluster and the centroid (mean synergy in the cluster). The clustering procedure was performed for both spatial and temporal model and for each *R*^*2*^ threshold.

### 2.5 Outcome measures and statistics

The goodness of the reconstruction was obtained from the original and the surrogate dataset with a spatial and a temporal model. A linear mixed-effects model (McLean et al., 1991) was fitted for the *R*^*2*^ in order to investigate the effects of the use of the surrogate and original dataset. First, the data was tested for normality with the Kolmogorov-Smirnov test. Then, the *R*^*2*^ was modelled as the dependent variable with fixed effects for dataset type and order of extraction with interaction, considering the order of extraction as a categorical variable. Random intercepts were included for effects of subjects. The level of significance (*α*) was set 0.05. In order to compare the spatial and the temporal model, we proceeded as follows. First, we compared the goodness of reconstruction obtained from each model and a statistical analysis was conducted to evaluate the differences. We extracted synergies according to 3 reconstruction *R*^*2*^ thresholds and we quantified the mean and the standard deviation of each order of factorization to compare the *R*^*2*^ achieved with spatial and temporal synergies. Linear mixed-effects model was fitted for the *R*^*2*^ to investigate the effects of the use of the spatial and temporal model. First, the data was tested for normality with the Kolmogorov-Smirnov test. Then, the *R*^*2*^ was modelled as the dependent variable with fixed effects for synergy model and order of extraction with interaction, considering the order of extraction as a categorical variable. Random intercepts were included for effects of subjects. The level of significance (*α*) was set 0.05.

Furthermore, we clustered extracted synergies across participants and *R*^*2*^ levels to define the inter-subject repertoire of invariant spatial and temporal synergies. The inter-subject similarity was computed via cosine angle, comparing all combination of synergies present in a cluster and then mediated for each level of *R*^*2*^. This process was done for each dataset and for both spatial and temporal invariant synergies.

Finally, to provide a detailed characterization of the temporal model and of the temporal structure found in the data, temporal synergies were extracted from the surrogate datasets, considering the same order of factorization of the original dataset for each *R*^*2*^ threshold, and they were clustered with k-means clustering algorithm. For both the original and the surrogate datasets, the mean of the temporal synergies in each cluster were fitted with a Gaussian distribution and the phase (the peak location), the period (distance between two consecutive phases), and the full width at half maximum (FWHM) were computed. After testing the data for normality with the Kolmogorov-Smirnov test, a two-sample t-test was performed at each *R*^*2*^ threshold in order to identify the difference between original and surrogate datasets. The level of significance (*α*) was set 0.05.

## 3. Results

### 3.1 Validation of the synergy models

In Figure 4, the comparison of the reconstruction *R*^*2*^ of the spatial model using the original and the surrogate data is shown for both REACH PLUS and NINAPRO dataset. The *R*^*2*^ reconstruction was higher using the original dataset with respect to the surrogate one, for both REACH PLUS and NINAPRO. For both datasets, the ‘dataset type’ (surrogate/original) had a significant influence on the reconstruction *R*^*2*^ (p < 0.001), indicating that the spatial organization of the muscle patterns captured by the spatial synergy decomposition did not simply arise from the amplitude distribution of the data.

**Figure 4.**
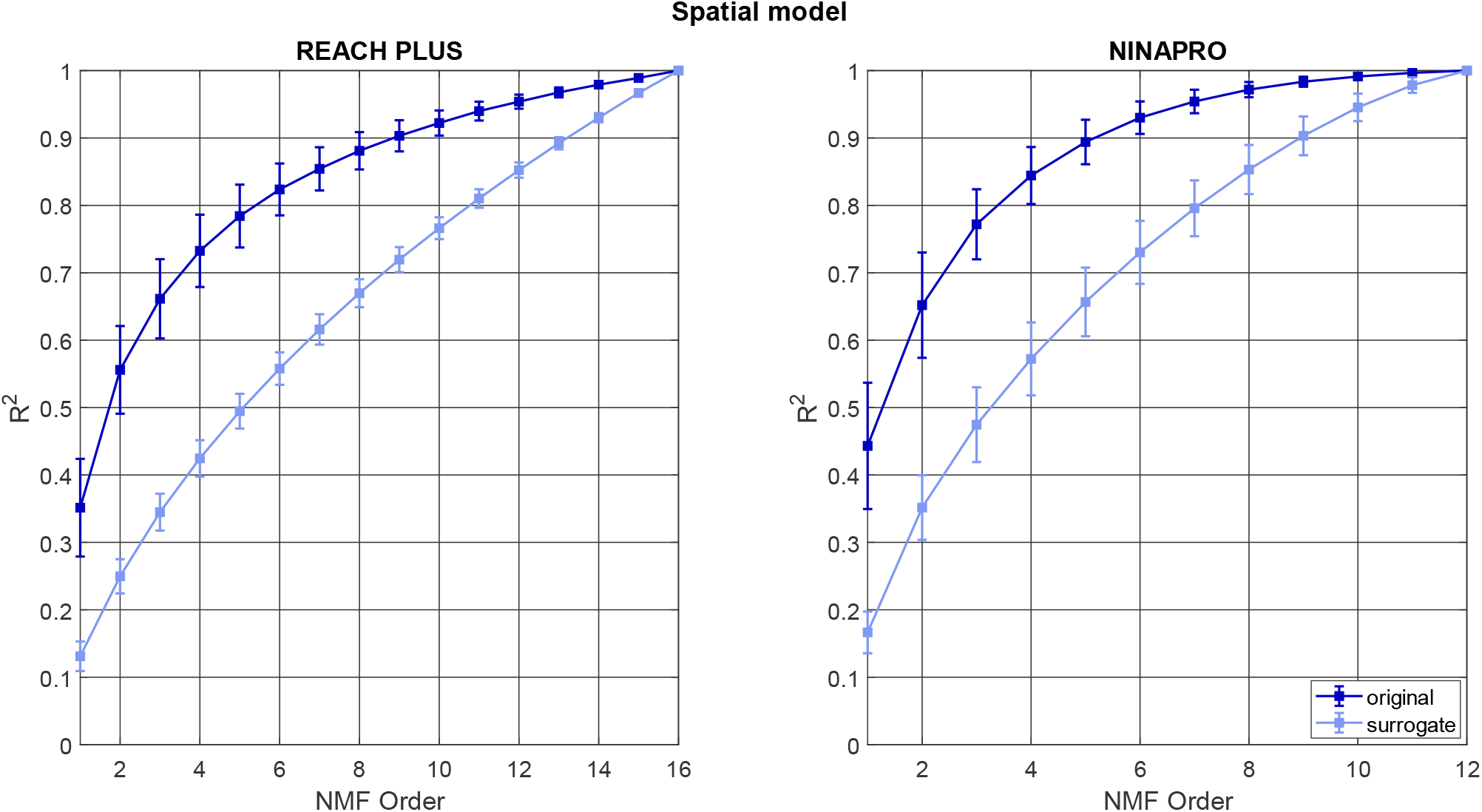
Reconstruction R^2^ computed with spatial synergy model: comparison between original data (dark blue) and the surrogate dataset (light blue). Both REACH PLUS and NINAPRO results are reported, averaged across participants. The squares are the mean across participants and the error bars represent the standard deviations.

The difference between the *R*^*2*^ curve of the surrogate and the *R*^*2*^ curve of the original data was significant for all the order of extraction with p < 0.001 until order 14 (included) for REACH PLUS and order 10 (included) for NINAPRO.

In Figure 5, the comparison of the reconstruction *R*^*2*^ of the temporal model using the original and the surrogate data is shown for both REACH PLUS and NINAPRO dataset. As for the spatial model, the reconstruction *R*^*2*^ was higher using the original dataset with respect to the surrogate one, for both REACH PLUS and NINAPRO. For both datasets, the ‘dataset type’ (surrogate/original) exhibited a significant influence on the reconstruction *R*^*2*^ (p < 0.001), indicating that also the temporal organization of the muscle patterns revealed by the temporal synergy decomposition could not simply be explained by the smoothness of the data.

**Figure 5.**
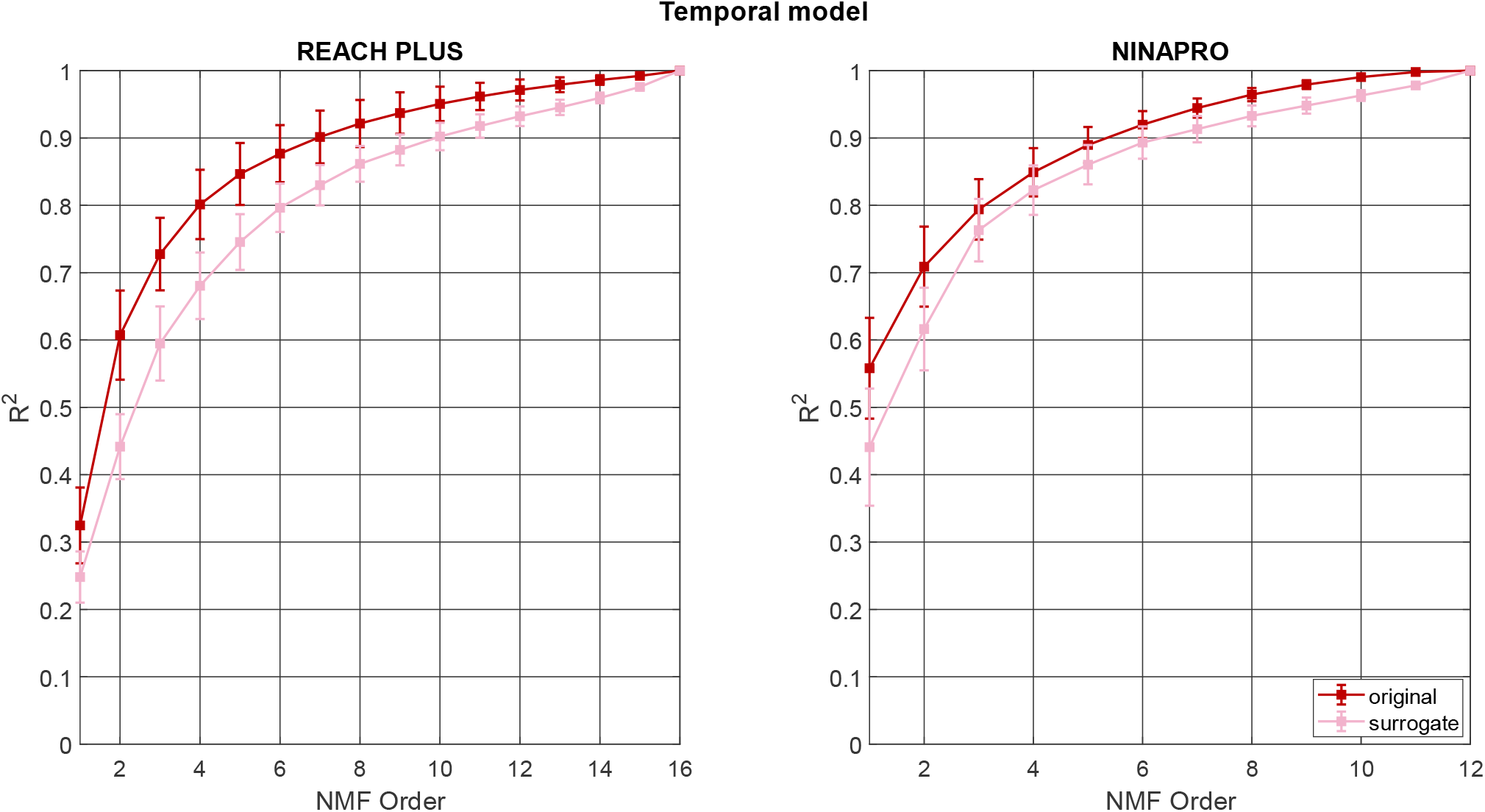
Reconstruction R^2^ computed with temporal synergy model: comparison between original data (red) and the surrogate dataset (pink). Both REACH PLUS and NINAPRO results are reported, averaged across participants. The squares are the mean across participants and the error bars represent the standard deviation.

The difference between the *R*^*2*^ curve of the surrogate and the *R*^*2*^ curve of the original data was significant for all the order of extraction until order 14 (p = 0.015) for REACH PLUS and order 11 (p = 0.03) for NINAPRO.

### 3.2 Synergy reconstruction and order of factorization

In Figure 6, we reported the reconstruction R^2^ comparing spatial synergies and temporal synergies for the REACH PLUS and for NINAPRO dataset.

**Figure 6.**
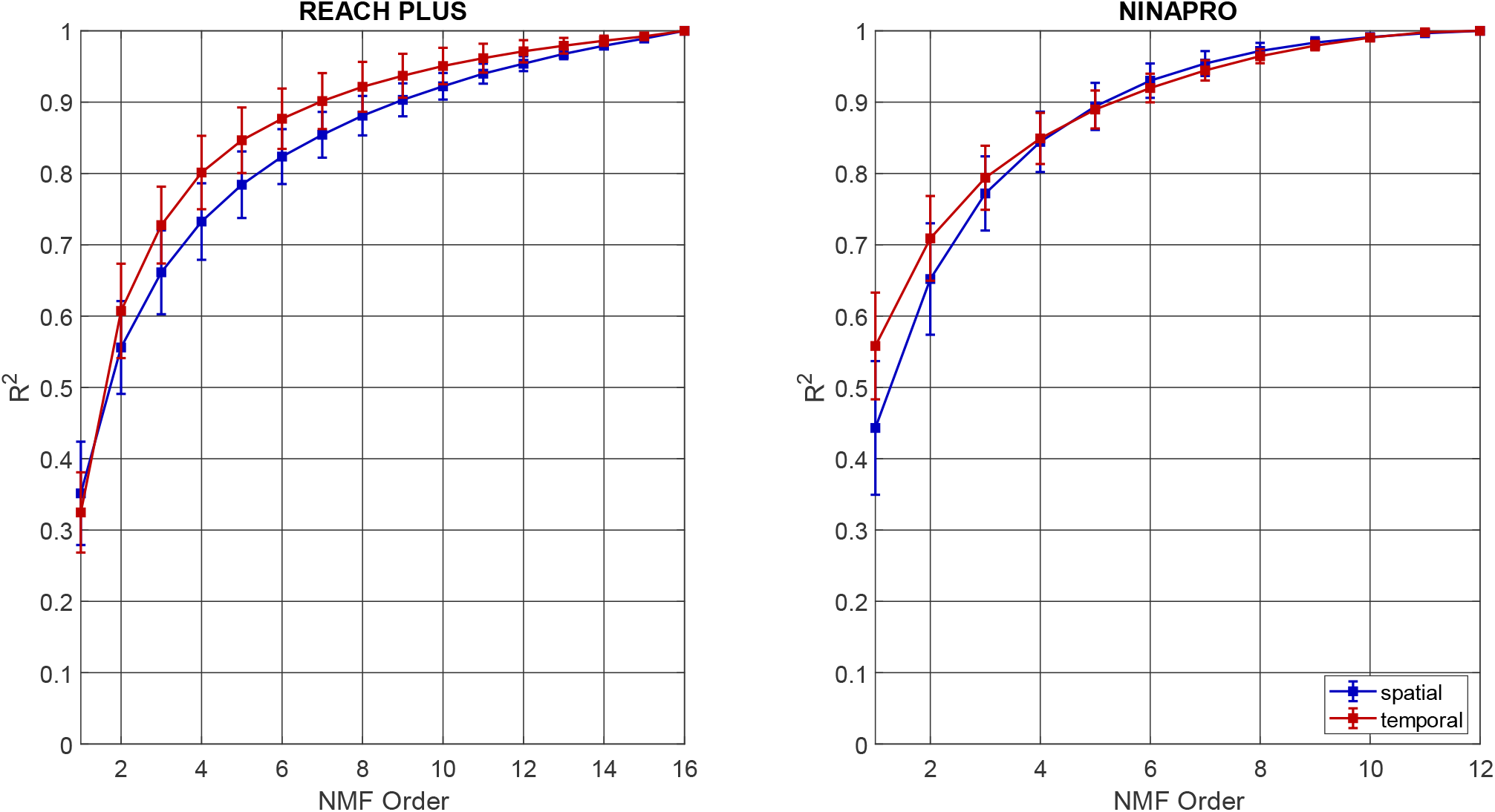
Reconstruction R^2^ for spatial and temporal synergies for both datasets (REACH PLUS and NINAPRO)

The *R*^*2*^ reconstruction was reported from order 1 to the number of EMG channels, 16 for REACH PLUS and 12 for NINAPRO. In REACH PLUS dataset, the two methods reported nearly the same *R*^*2*^ at order 1 (0.35 for spatial and at 0.32 for temporal). For all the other order of factorization, the temporal model showed a higher *R*^*2*^. In NINAPRO dataset, instead, the *R*^*2*^ curves of the two methods crossed each other between order 4 and 5: *R*^*2*^ is higher in temporal synergies until order 4 and in spatial synergies after this order. The linear mixed-effects model analysis indicated that the ‘synergy model’ had significant effects on the *R*^*2*^, with p < 0.001 for REACH PLUS and with p = 0.04 for NINAPRO, indicating a different reconstruction between the spatial and the temporal model. Furthermore, for REACH PLUS the difference between the *R*^*2*^ curve of the spatial model and the *R*^*2*^ curve of the temporal model was significant for all the order of extraction until order 10 (p = 0.028), while for NINAPRO the difference between the *R*^*2*^ of the two models is significant only for order 1 and 2 with p < 0.001.

In table I, the orders of factorization averaged across participants for each *R*^*2*^ threshold are reported. For the REACH PLUS dataset, the number of extracted synergies for each *R*^*2*^ thresholds varied across participants respectively from 4 to 7, from 5 to 9, from 7 to 11 for the spatial model, while the range was between 3 and 6, 4 and 7, 5 and 10, for temporal model. For NINAPRO dataset, the order of factorization ranged from 2 to 5, from 3 to 6, from 3 to 8 for the spatial model, while using temporal model the orders were between 2 and 5, 3 and 6, 4 and 8.

**Table I.** Means and standard deviations for the number of extracted synergies for each R^2^ threshold for spatial and temporal models in both datasets.

| MODEL | $R^2 = 0.80$ | $R^2 = 0.85$ | $R^2 = 0.90$ |
| --- | --- | --- | --- |
| **REACH PLUS** |  |  |  |
| SPATIAL | $5.8 \pm 1.0$ | $7.2 \pm 1.2$ | $9.2 \pm 1.2$ |
| TEMPORAL | $4.6 \pm 0.8$ | $5.7 \pm 1.2$ | $7.5 \pm 1.7$ |
| **NINAPRO** |  |  |  |
| SPATIAL | $3.8 \pm 0.7$ | $4.6 \pm 0.8$ | $5.6 \pm 0.9$ |
| TEMPORAL | $3.6 \pm 0.8$ | $4.3 \pm 0.8$ | $5.7 \pm 0.8$ |

In Figure 7, we report an example of spatial synergies and variant temporal coefficients extracted from EMG signal of a typical subject of the REACH PLUS dataset. With *R*^*2*^ threshold = 0.80, the order of extraction was five, with *R*^*2*^ = 0.81. In the first row, the distribution of muscles in each synergy is shown, while the temporal activation profiles of each synergy are reported for each direction of movement.

**Figure 7.**
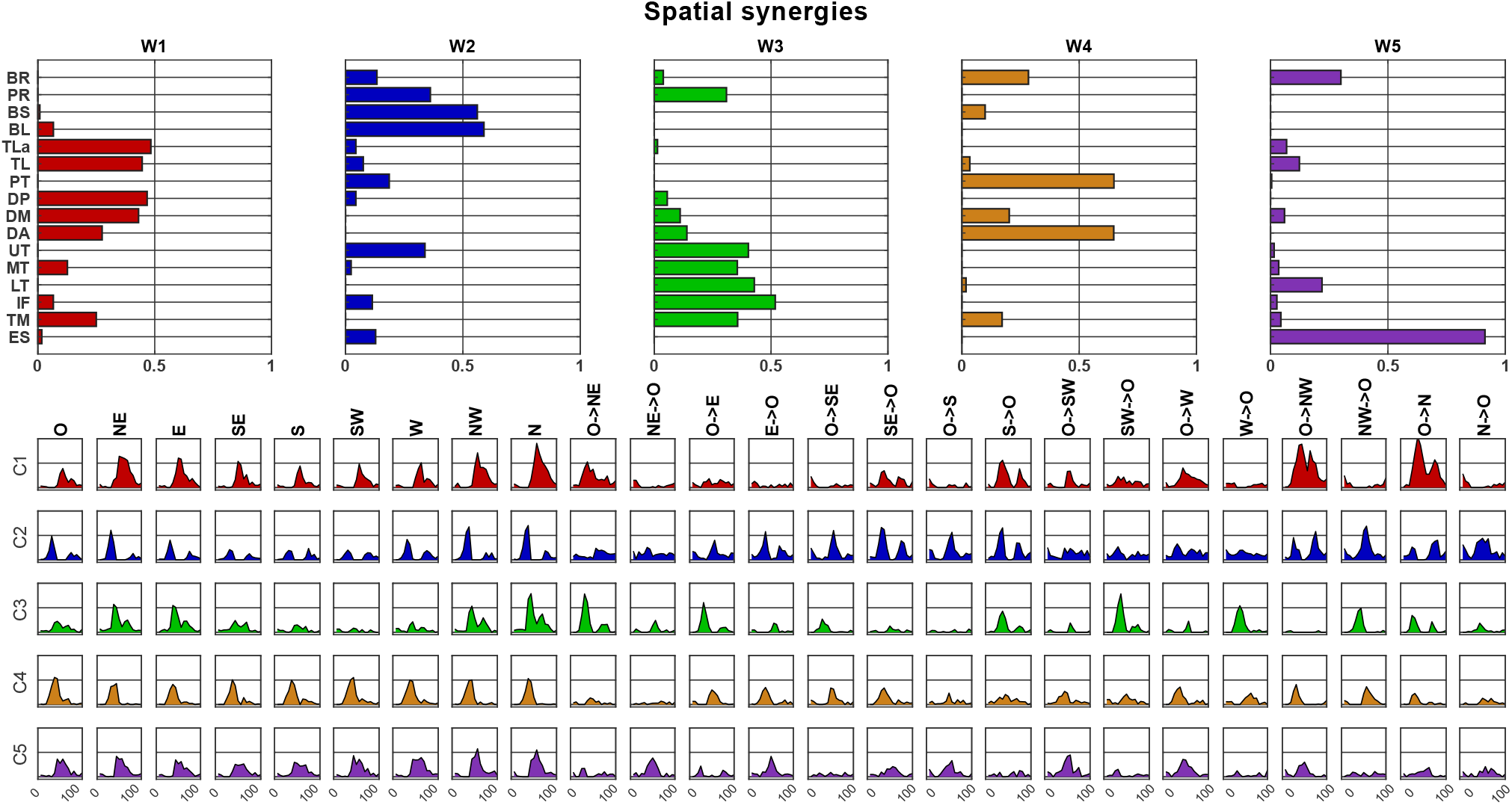
Typical example of spatial synergies and variant temporal coefficients, extracted from the REACH PLUS dataset.

An example of temporal synergies with variant spatial loads extracted from the same participant is reported in Figure 7. In this case, imposing a *R*^*2*^ threshold of 0.80, four synergies were extracted, (*R*^*2*^ = 0.83). The invariant temporal synergies are shown in the first column and the respective variant spatial loads are reported for each direction of movement.

### 3.3 Inter-individual variability of invariant components

In Figure 9, the spatial synergies after clustering are reported for the REACH PLUS dataset (inter-subject synergies).

**Figure 8.**
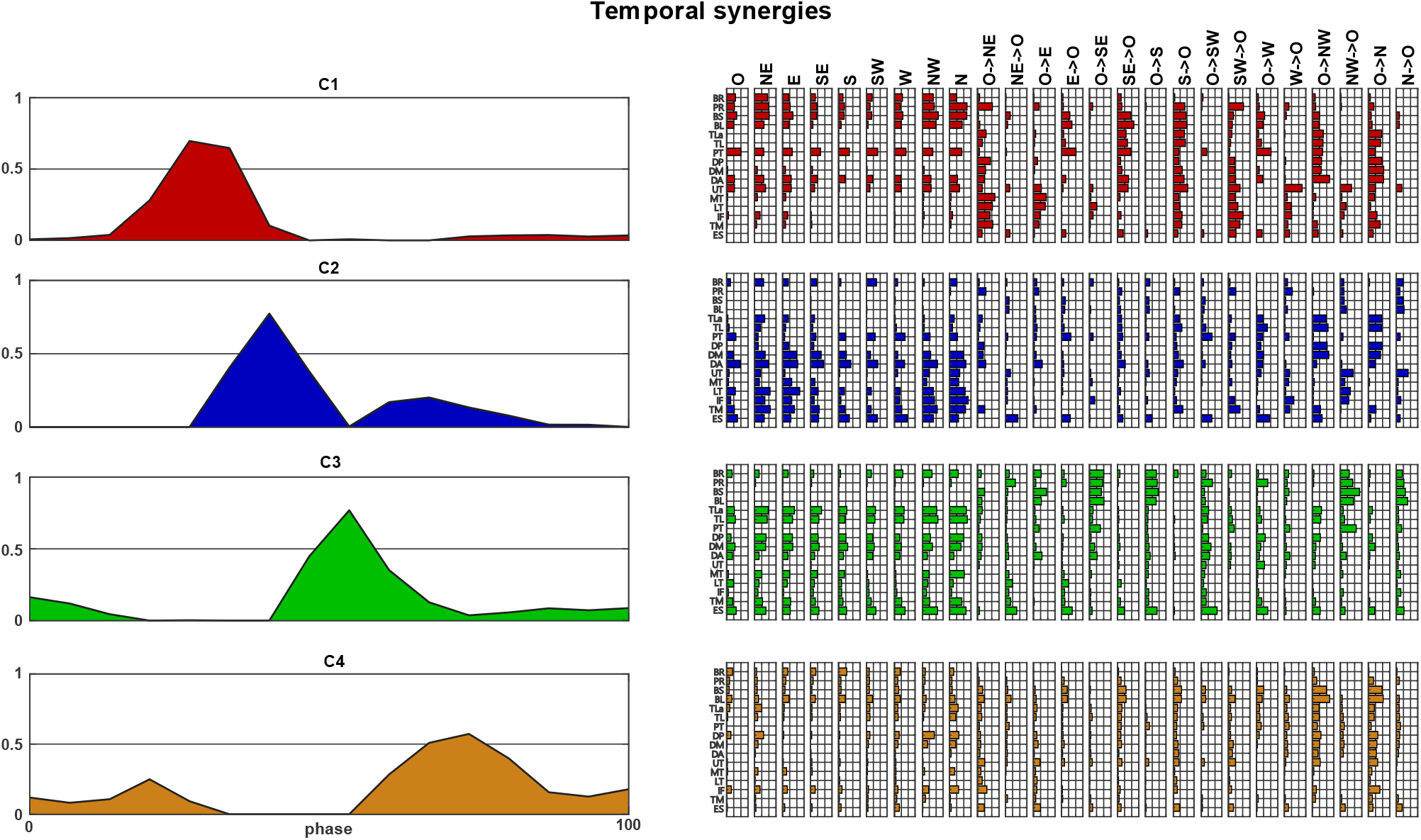
Typical example of temporal synergies and variant spatial loads, extracted from the REACH PLUS dataset.

**Figure 9.**
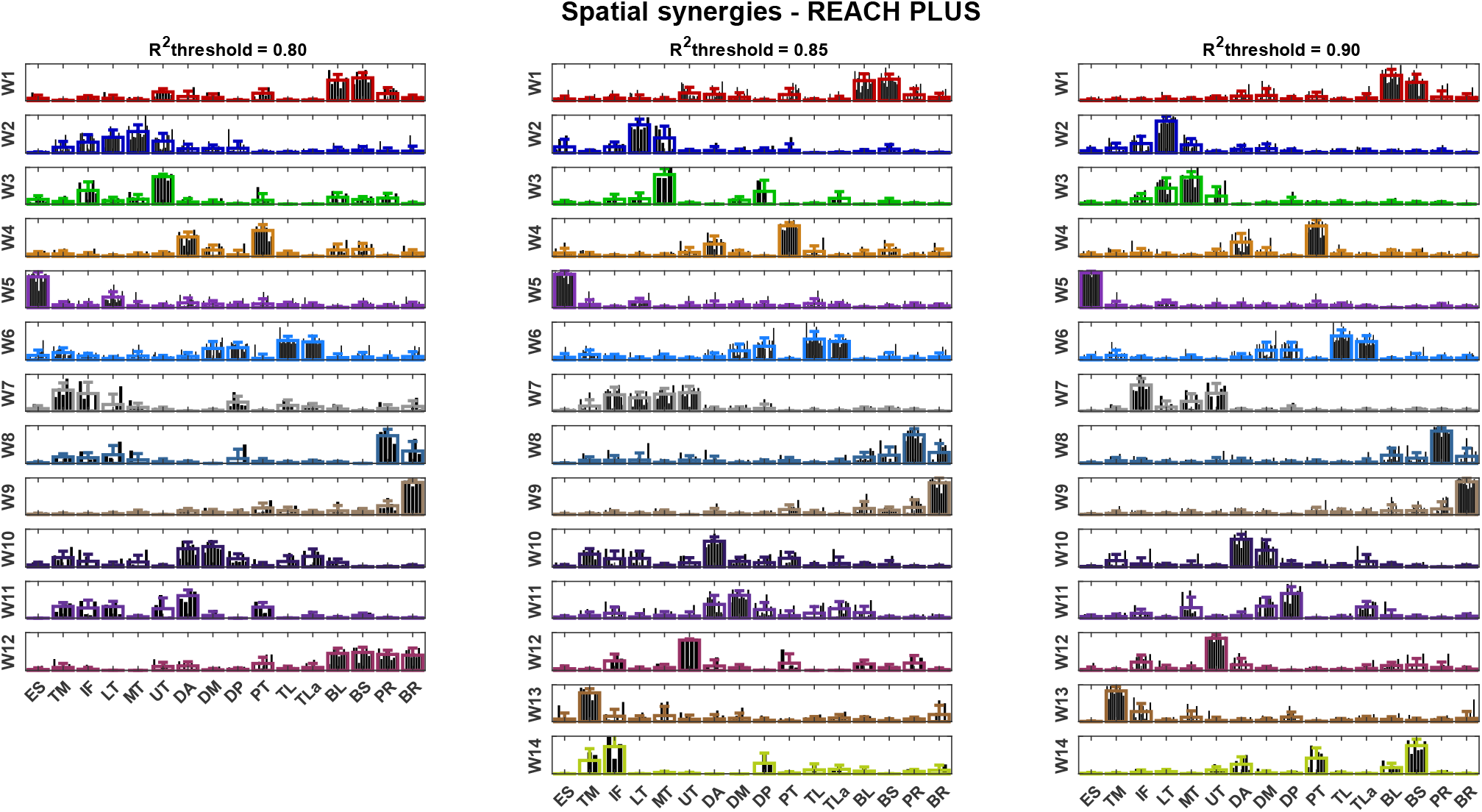
Spatial synergies averaged across participants after k-means clustering, for three level of reconstruction R^2^ (REACH PLUS). The bold lines represent the means and the standard deviations of the synergies for each cluster.

Twelve clusters were chosen for grouping the synergies obtained with *R*^*2*^ = 0.80 and each cluster contained from 4 to 14 synergies. With *R*^*2*^ = 0.85, fourteen clusters were found to group synergies, with 3 to 14 synergies in each cluster. Finally, for *R*^*2*^ = 0.90, synergies were clustered in fourteen groups composed of 6 to 16 synergies each.

In Figure 10, we report the temporal synergies after matching across participants for REACH PLUS (inter-subject synergies).

**Figure 10.**
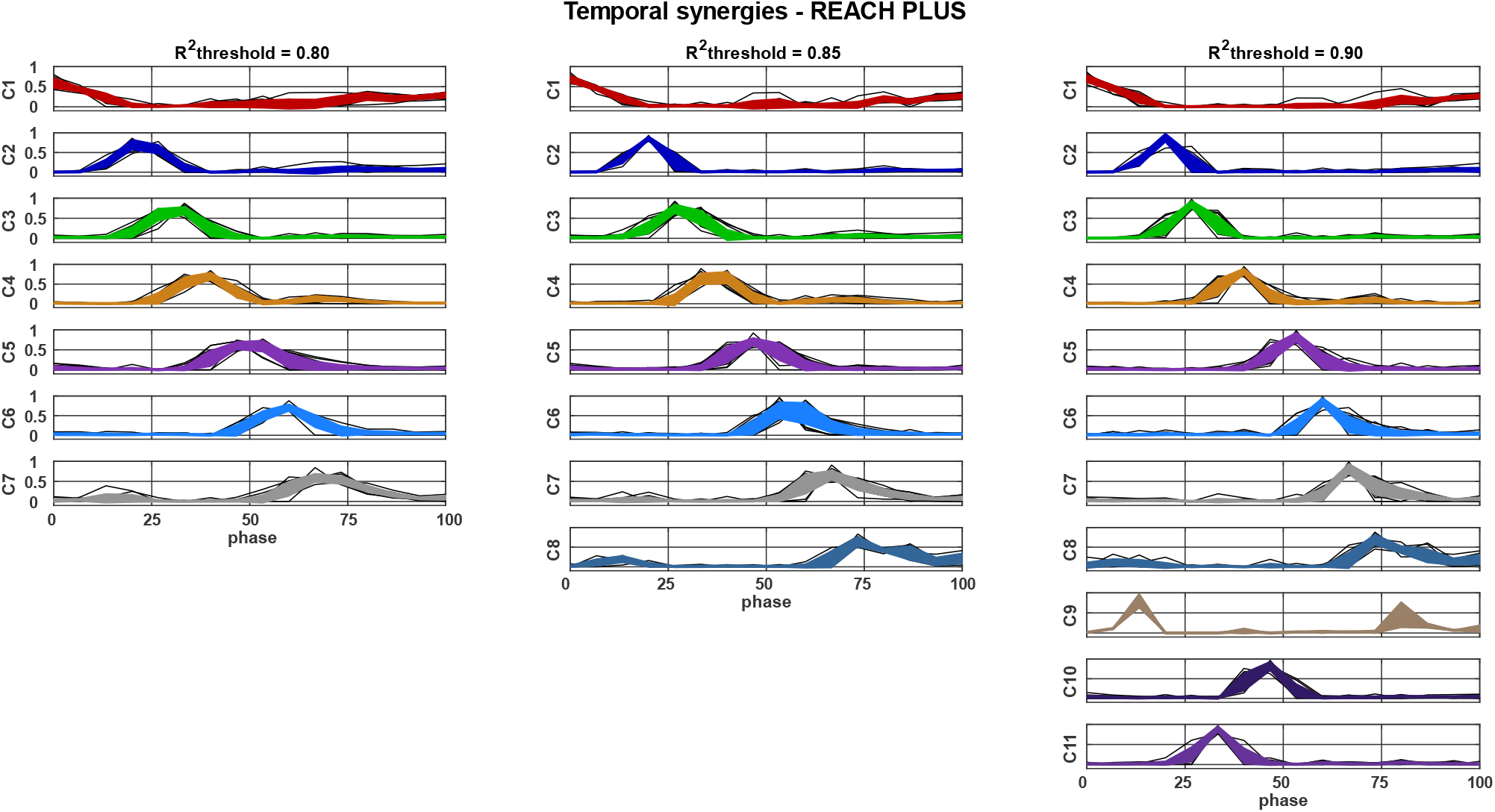
Temporal synergies averaged across participants after k-means clustering, for three level of reconstruction R^2^ for REACH PLUS dataset. The bold lines represent the means with the standard deviations of the synergies for each cluster.

For the first *R*^*2*^ threshold, temporal synergies were clustered in seven groups, containing from 8 to 14 synergies. With *R*^*2*^ = 0.85, eight clusters were used to match the temporal synergies and in each cluster the number of synergies ranged from 6 to 15. Eleven clusters were found for *R*^*2*^ = 0.90, containing from 2 to 14 synergies each. The number of clusters needed to group temporal synergies were lower with respect to spatial synergies for all the *R*^*2*^ thresholds, as reflected by the lower mean order of factorization.

In Figure 11, we report the spatial synergies after matching across participants for the NINAPRO dataset (inter-subject synergies).

**Figure 11.**
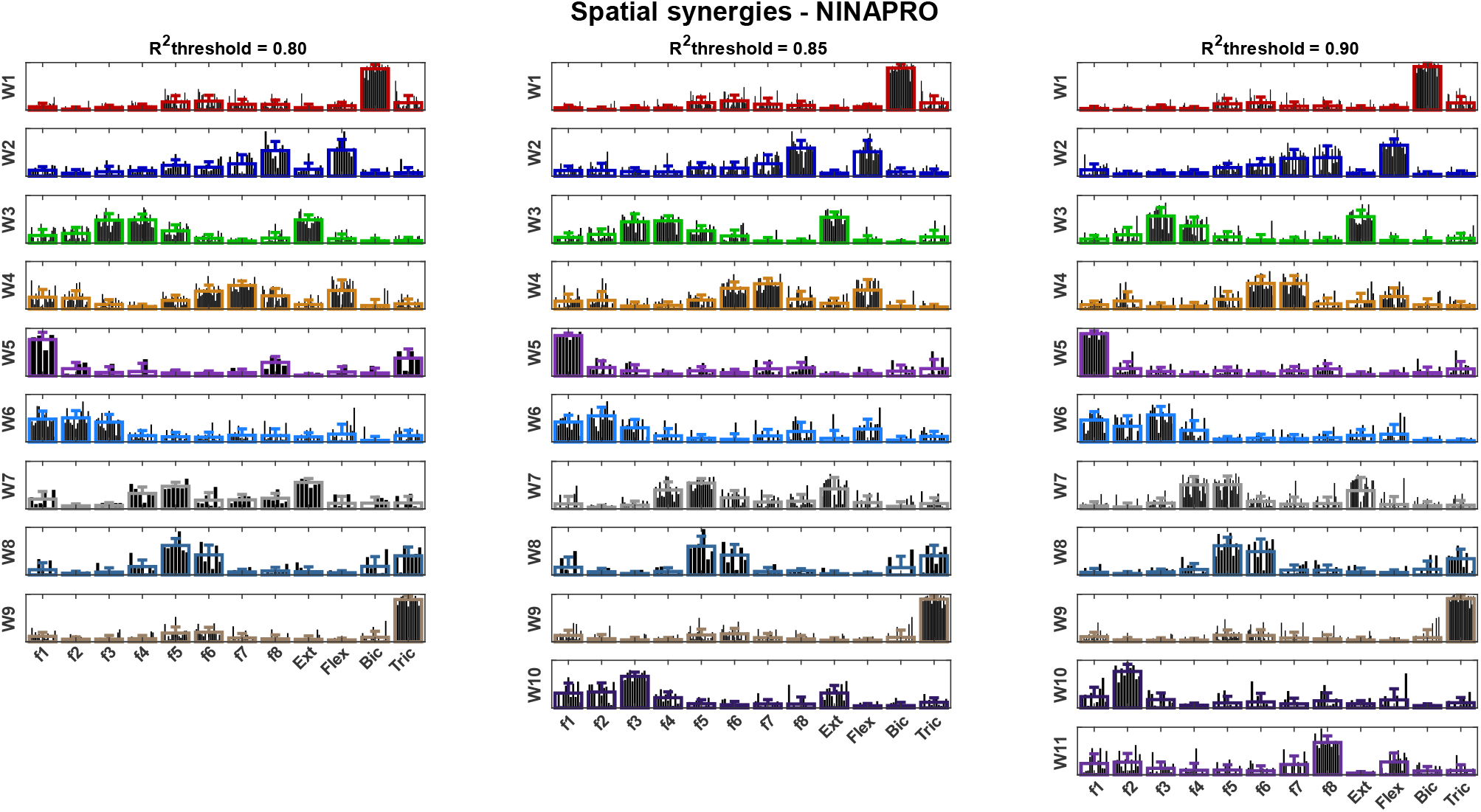
Spatial synergies averaged across participants after clustering procedure, at three level of reconstruction R^2^ for NINAPRO dataset. The bold lines represent the means with the standard deviations of the synergies belonging to that cluster.

For the NINAPRO dataset, the number of clusters for *R*^*2*^ = 0.80 was nine, with a range of 4 to 23 synergies in each group. For *R*^*2*^ = 0.85, spatial synergies were clustered in ten groups (each composed of 6 to 24 synergies). Finally, for *R*^*2*^ = 0.90, eleven clusters were found and each cluster contained from 9 to 25 synergies.

In Figure 12, we report the temporal synergies after matching across participants for the NINAPRO dataset (inter-subject synergies).

**Figure 12.**
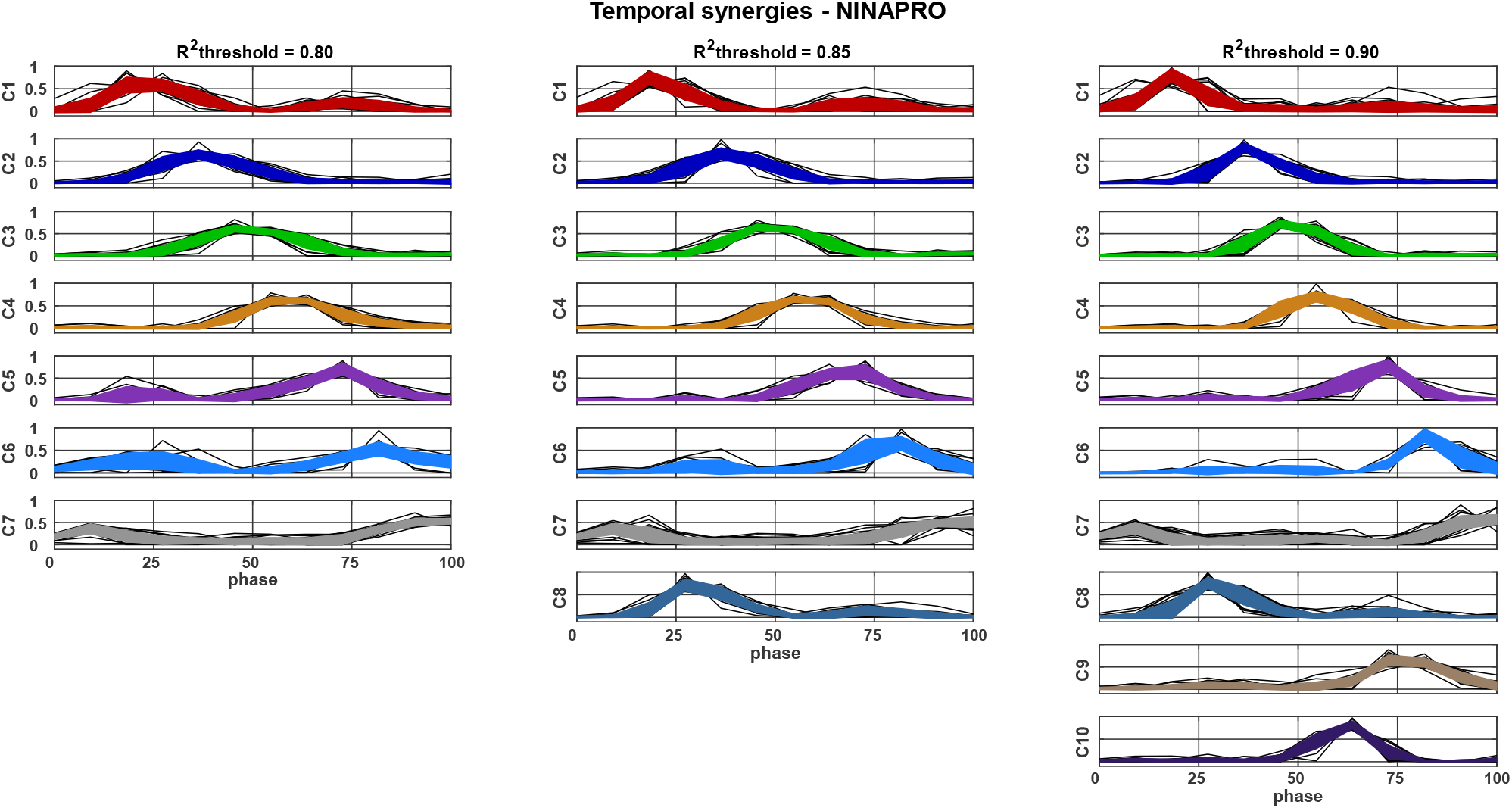
Temporal synergies averaged across participants after k-means clustering, for three level of reconstruction R^2^ for NINAPRO dataset. The bold lines represent the means with the standard deviations of the synergies for each cluster.

Temporal synergies were grouped in seven clusters for *R*^*2*^ = 0.80, with 10 to 21 synergies in each group. For *R*^*2*^ equal to 0.85, the number of clusters was eight and each one contained from 12 to 27 temporal synergies. For *R*^*2*^ = 0.90, ten clusters were found and the number of synergies in each one ranged from 10 to 27. As for REACH PLUS dataset, the number of clusters were lower for the temporal model with respect to the spatial model in all the *R*^*2*^ thresholds, even if the mean order of extraction was similar between the two models.

In Table II, the inter-individual similarities for spatial and temporal synergies in the same cluster are summarized.

**Table II.** Inter-individual similarity (spatial and temporal synergies) is reported for each threshold and each dataset. The values are the mean and standard deviations between the similarity computed in each cluster obtained at that threshold.

| MODEL | $R^2 = 0.80$ | $R^2 = 0.85$ | $R^2 = 0.90$ |
| --- | --- | --- | --- |
| **REACH PLUS** |  |  |  |
| SPATIAL | $0.78 \pm 0.04$ | $0.77 \pm 0.04$ | $0.81 \pm 0.04$ |
| TEMPORAL | $0.89 \pm 0.02$ | $0.88 \pm 0.02$ | $0.89 \pm 0.02$ |
| **NINAPRO** |  |  |  |
| SPATIAL | $0.85 \pm 0.03$ | $0.86 \pm 0.02$ | $0.86 \pm 0.03$ |
| TEMPORAL | $0.90 \pm 0.03$ | $0.89 \pm 0.03$ | $0.88 \pm 0.02$ |

The similarity computed in each cluster was high (>0.75) for all the conditions. In the REACH PLUS dataset, similarity of spatial synergies was 0.78, 0.77 and 0.81 for the 0.80, 0.85, 0.90 thresholds, respectively. For the temporal model, instead, the similarity was higher, reaching 0.89 for *R*^*2*^ threshold = 0.80 and 0.90 and 0.88 for *R*^*2*^ threshold = 0.85. For the NINAPRO dataset, the similarity of spatial synergies was 0.85, 0.86 and 0.86. For temporal synergies, the similarity was higher and decreased when increasing the threshold: it was 0.90 at *R*^*2*^ threshold = 0.80, 0.89 for *R*^*2*^ threshold = 0.85 and 0.88 for the highest threshold.

Similarity was computed in each cluster and, then, mediated across clusters for the same *R*^*2*^ threshold. Mean and standard deviations are compared in Figure 13. For both the datasets, temporal synergies showed a higher similarity and lower standard deviations with respect to spatial synergies.

**Figure 13:**
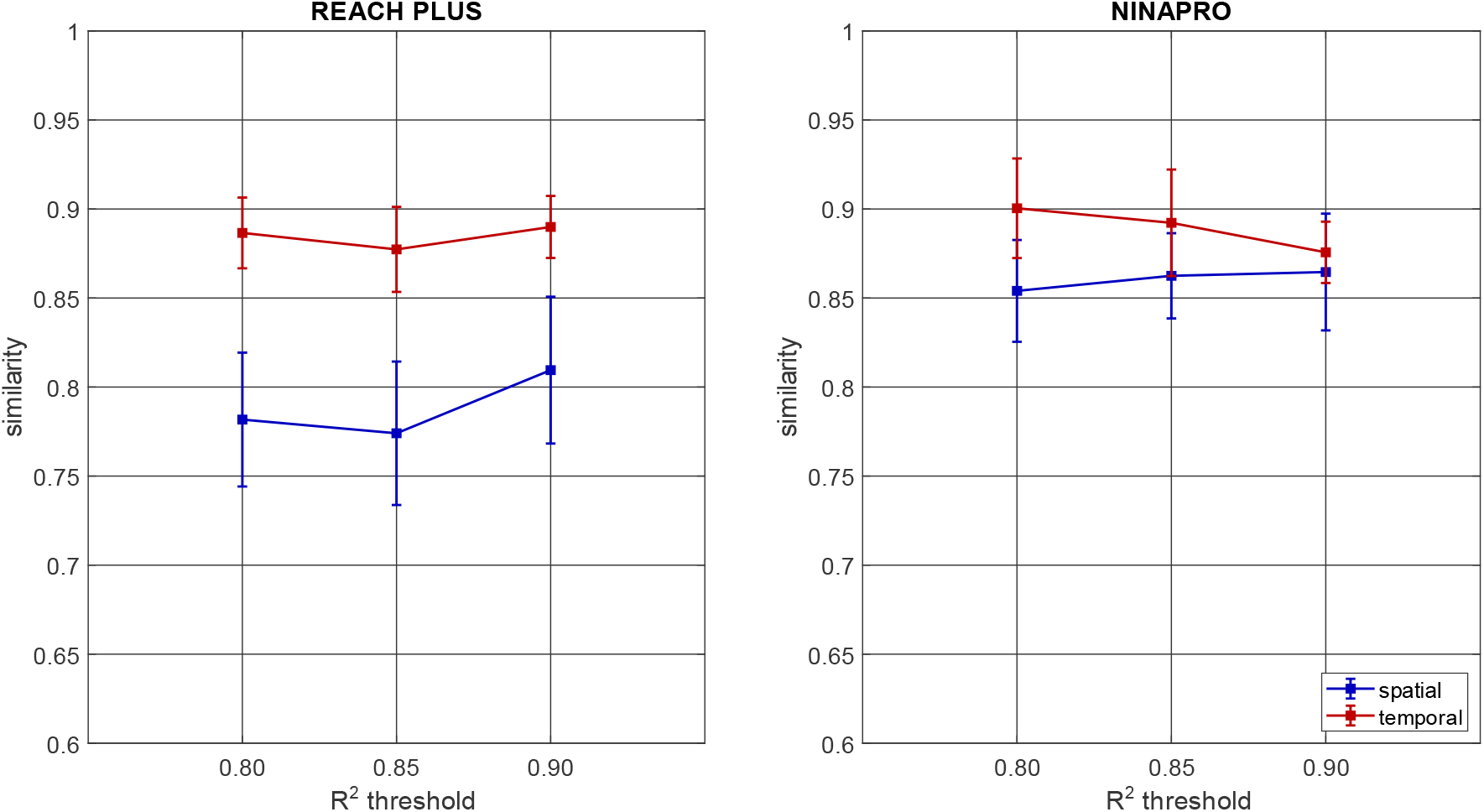
Inter-subject similarity of invariant synergies (spatial and temporal) for both datasets when changing the R^2^ threshold.

### 3.4 Characterization of the temporal synergies

In Figure 14, the results of the k-means clustering on the temporal synergies extracted from the surrogate data of REACH PLUS dataset are reported. For each subject, the order of factorization for each threshold was the same used for the synergy extraction from the original dataset and the extracted synergies were clustered in 7, 9, and 11 clusters. The number of clusters was equal to the number of clusters of the original dataset for *R*^*2*^ threshold = 0.80 and *R*^*2*^ threshold = 0.90, while for *R*^*2*^ threshold = 0.85 the number of clusters was higher.

**Figure 14.**
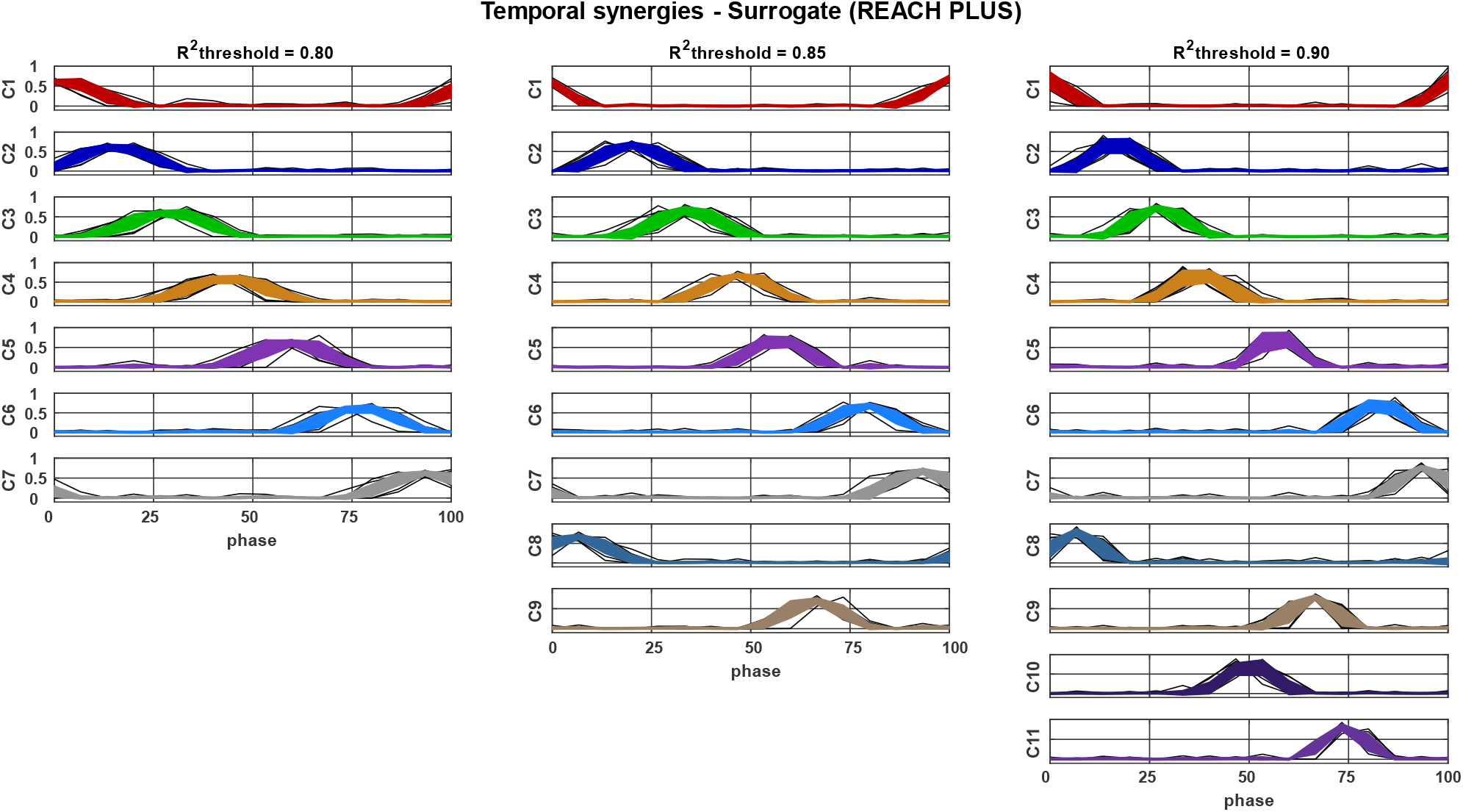
Temporal synergies extracted from surrogate dataset averaged across participants after k-means clustering, for three level of reconstruction R^2^ for the REACH PLUS dataset. The bold lines represent the means with the standard deviations of the synergies for each cluster.

In Figure 15, the results of the clustering procedure on the temporal synergies extracted from the surrogate data of NINAPRO dataset are reported. For each subject, the order of factorization for each threshold was the same used for the synergy extraction from the original dataset and the extracted synergies were clustered in 6, 7, and 10 clusters. Differently from the REACH PLUS dataset, the clusters were fewer than the clusters of the original dataset *R*^*2*^ threshold = 0.80 and *R*^*2*^ threshold = 0.85, indicating that the synergies extracted from the surrogate dataset were more similar between subjects, while for *R*^*2*^ threshold = 0.90 the number of clusters was the same of the original dataset.

**Figure 15.**
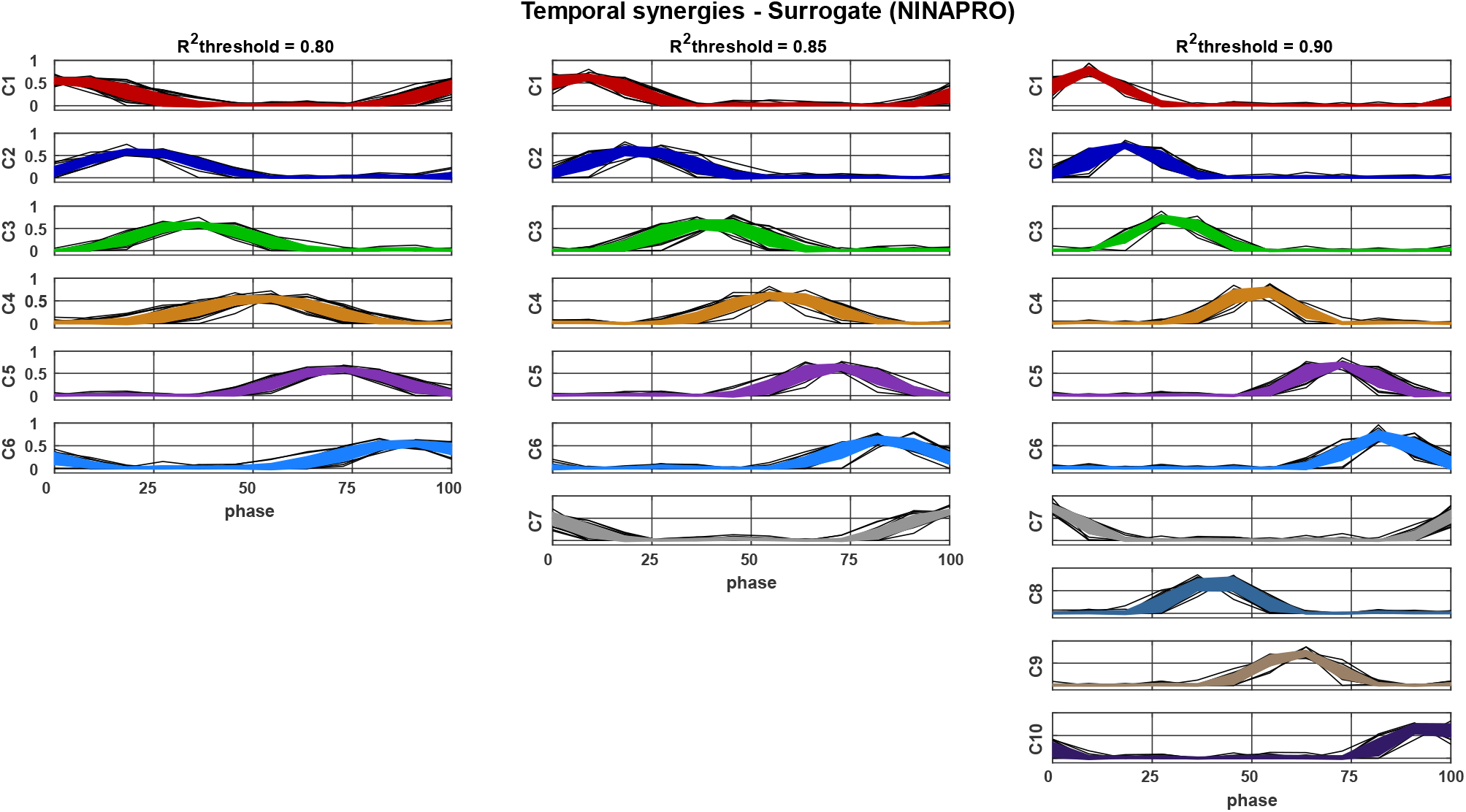
Temporal synergies extracted from surrogate dataset averaged across participants after k-means clustering, at three level of reconstruction R^2^ for NINAPRO dataset. The bold lines represent the means with the standard deviations of the synergies for each cluster.

The mean temporal synergy of each cluster was fitted with a Gaussian function and the phase of the peak and its full width at half maximum (FWHM) were computed for both the original and the surrogate dataset and they are reported in Table III. The synergies with normalized movement phase <5% or >95% were reported in brackets and excluded from the computation of the mean period (difference between peak phases) and FWHM because they represent the tails of the signals and these curves were incomplete and cannot be easily fitted with a Gaussian function. Mean widths and mean periods are reported in Figure 16.

**Table III.** Full widths at half maximum and phases of the peak are reported for the mean temporal synergy of each cluster of the original (Or) and the surrogate (Sur) dataset of both REACH PLUS and NINAPRO datasets. In the last column, the mean width and the mean phase difference between consecutive peaks (period) are reported. The values in brackets are excluded from the mean since the phases was <5 or >95. The bold indicates the synergies in which the FWHM of the original data is larger than the FWHM of the surrogate data.

| REACH PLUS |  |  |  |  |  |  |  |  |  |  |  |  |  |  |
| --- | --- | --- | --- | --- | --- | --- | --- | --- | --- | --- | --- | --- | --- | --- |
|  |  | R<sup>2</sup> | C1 | C2 | C3 | C4 | C5 | C6 | C7 | C8 | C9 | C10 | C11 | mean |
| FWHM | Or | 0.80 | (23.1) | 13.5 | 14.2 | 14.7 | 17.5 | 16.1 | **21.9** |  |  |  |  | 16.3±3.1 |
|  |  | 0.85 | (21.6) | 10.7 | 13.5 | 12.4 | 15.3 | 15.5 | **19.4** | **21.7** |  |  |  | 15.5±3.9 |
|  |  | 0.90 | (21.3) | 9.9 | 10.9 | 10.9 | 11.3 | 10.0 | 10.7 | **16.9** | 5.1 | 10.8 | 9.7 | 10.6±2.8 |
|  | Sur | 0.80 | (19.1) | 19.7 | 20.8 | 21.4 | 19.8 | 19.3 | 20.6 |  |  |  |  | 20.3±0.8 |
|  |  | 0.85 | (8.0) | 16.9 | 17.1 | 17.2 | 15.5 | 16.7 | 17.3 | 17.4 | 15.8 |  |  | 16.7±0.7 |
|  |  | 0.90 | (9.1) | 13.2 | 14.5 | 14.4 | 11.1 | 13.1 | 13.7 | 12.3 | 12.4 | 12.9 | 12.1 | 13.0±1.1 |
| phase | Or | 0.80 | (-0.9) | 22.4 | 30.8 | 38.2 | 49.7 | 59.1 | 70.4 |  |  |  |  | 9.6±1.8 |
|  |  | 0.85 | (-1.5) | 19.7 | 28.2 | 37.0 | 47.3 | 56.5 | 67.4 | 77.8 |  |  |  | 9.7±1.0 |
|  |  | 0.90 | (-1.6) | 19.7 | 27.1 | 38.8 | 52.4 | 60.3 | 67.8 | 76.4 | 11.4 | 45.5 | 33.1 | 7.2±1.0 |
|  | Sur | 0.80 | (2.0) | 15.2 | 28.6 | 43.5 | 59.5 | 77.0 | 93.5 |  |  |  |  | 15.7±1.5 |
|  |  | 0.85 | (1.0) | 20.0 | 34.2 | 47.2 | 56.7 | 78.6 | 93.0 | 5.9 | 66.0 |  |  | 12.4±2.2 |
|  |  | 0.90 | (1.0) | 16.8 | 27.2 | 37.3 | 57.1 | 82.6 | 93.9 | 6.5 | 66.1 | 50.2 | 74.0 | 9.7±1.8 |

| NINAPRO |  |  |  |  |  |  |  |  |  |  |  |  |  |  |
| --- | --- | --- | --- | --- | --- | --- | --- | --- | --- | --- | --- | --- | --- | --- |
|  |  | R<sup>2</sup> | C1 | C2 | C3 | C4 | C5 | C6 | C7 | C8 | C9 | C10 | C11 | mean |
| FWHM | Or | 0.80 | 22.5 | 26.9 | 28.5 | 25.4 | 22.9 | 31.8 | (38.8) |  |  |  |  | 26.3±3.5 |
|  |  | 0.85 | 16.5 | 24.6 | 24.5 | 23.9 | 23.8 | 22.0 | (42.7) | 18.8 |  |  |  | 22.0±3.2 |
|  |  | 0.90 | 14.8 | 16.5 | 19.5 | **21.5** | 17.4 | 14.4 | (25.2) | 16.1 | **22.0** | 15.8 |  | 17.5±2.8 |
|  | Sur | 0.80 | (33.1) | 30.9 | 30.4 | 34.8 | 30.5 | 35.7 |  |  |  |  |  | 32.5±2.6 |
|  |  | 0.85 | 26.5 | 27.1 | 27.2 | 27.5 | 25.2 | 26.7 | (25.4) |  |  |  |  | 26.5±1.0 |
|  |  | 0.90 | 18.6 | 20.2 | 19.4 | 19.5 | 21.2 | 20.4 | (13.9) | 20.0 | 19.9 | 20.7 |  | 19.5±2.2 |
| phase | Or | 0.80 | 24.9 | 37.5 | 49.0 | 59.8 | 71.3 | 82.6 | (98.0) |  |  |  |  | 11.7±1.0 |
|  |  | 0.85 | 20.3 | 38.4 | 49.6 | 57.6 | 67.8 | 79.6 | (99.5) | 30.8 |  |  |  | 9.9±1.7 |
|  |  | 0.90 | 19.1 | 37.9 | 48.3 | 55.2 | 70.6 | 82.9 | (97.2) | 29.7 | 77.6 | 61.8 |  | 8.0±1.9 |
|  | Sur | 0.80 | (2.7) | 21.5 | 34.9 | 51.7 | 71.5 | 89.1 |  |  |  |  |  | 16.9±2.7 |
|  |  | 0.85 | 7.4 | 21.5 | 39.3 | 56.5 | 71.3 | 83.7 | (100.3) |  |  |  |  | 15.3±2.2 |
|  |  | 0.90 | 8.7 | 18.2 | 29.8 | 51.2 | 71.4 | 83.1 | (101.7) | 41.0 | 61.1 | 94.9 |  | 10.8±0.9 |

**Figure 16.**
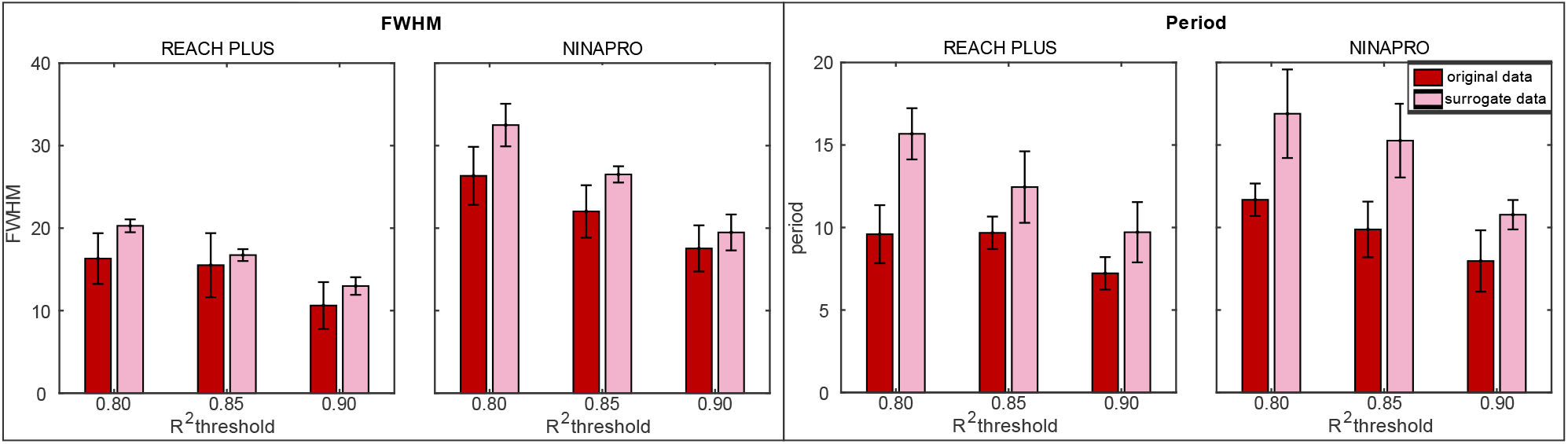
The mean FWHM (left panel) and the mean period (right panel) computed on the mean temporal synergy of each cluster are reported for both REACH PLUS and NINAPRO datasets. The red bars refer to the original data, while the pink ones to the surrogate data.

For both the original and surrogate datasets, the period and the FWHM decreased as the number of extracted synergies increased. The surrogate synergies showed larger FWHM and period with respect to the original synergies for all the thresholds, even when the number of clusters was higher (as in the REACH PLUS dataset). The original synergies appear more concentrated, with closer and narrower peaks.

Statistical analysis showed that FWHM was significantly different between original and surrogate data for *R*^*2*^ = 0.80 (p = 0.01) and *R*^*2*^ = 0.90 (p = 0.02) for REACH PLUS, while significant differences were found for *R*^*2*^ = 0.80 (p = 0.01) and *R*^*2*^ = 0.85 (p = 0.007) for NINAPRO. Differences in periods were statistically significant for all the threshold in both REACH PLUS and NINAPRO datasets.

## 4. Discussion

In this study, we performed a quantitative assessment of the spatial and temporal synergies extracted using NMF from two datasets that provide a comprehensive description of both distal and proximal upper limb movements. We compared spatial and temporal synergies in terms of goodness of reconstruction and inter-individual variability. We first showed that there exists a significative spatial and temporal structure in the EMG signals of several muscles that could not be simply explained by the amplitude distribution or the smoothness of the data. In fact, the goodness of reconstruction with temporal and spatial synergies obtained with a surrogate dataset when the temporal and spatial structures of individual muscles was randomized (while maintaining the same EMG spectral features) was lower than the one obtained with the original signals. Spatial and temporal synergy models were then compared in terms of quality of reconstruction (*R*^*2*^), number of synergies (reconstruction order) selected according to three different *R*^*2*^ thresholds, and inter-subject synergy variability. Interestingly, we found that especially for low and frequently used reconstruction *R*^*2*^, the temporal factorization required fewer temporal components than spatial ones. These findings were not analyzed in detail before and suggest that the smoothness of the data constrains the temporal dimensionality more than the amplitude distribution of the data constrains the spatial dimensionality. The differences between the two models were more evident in the proximal upper limb dataset. At the same time, the “elbow” in the reconstruction *R*^*2*^ curve was more pronounced with temporal synergies than with spatial synergies. It follows that it is often possible to find an even more compact representation of movement with temporal synergies with respect to the standard spatial model. It is indeed likely to obtain a lower dimensional representation when using the temporal model. On the contrary, spatial synergies were more parsimonious than temporal for higher reconstruction *R*^*2*^ values, that are less frequently used. Comparing both models with the surrogate dataset, the *R*^*2*^ curve of the surrogate data is closer to the *R*^*2*^ curve of the original data for the temporal model. Since the surrogate data indicate the intrinsic dimensionality expected for signal with such characteristics, the temporal synergies may have a lower dimensionality due to the smoothness of the data rather than to a more compact representation of the temporal model. Thus, the smoothness of the data appears to constrain the temporal dimensionality more than the amplitude distribution of the data constrains the spatial dimensionality. Consequently, according to this point of view, the spatial model gives a more compact representation of the movement, even if it requires more synergies as previously explained. However, we remark that spatial and temporal synergies capture different levels of organization and no assumption should be made a priori on the proper number of synergies that should be extracted (which can in general be different when adopting on model or the other). We conclude that the two methods are complementary at a mathematical level, even if they may reflect different features and organization at neural level. It follows that it is probably not correct to support only the use of the spatial (or the temporal) model.

The sensibility of the temporal synergy model on the number of samples of the input signals should be investigated more in future work. Decreasing the number of samples may give a higher reconstruction of the signal, as in Torricelli et al. (2020) in which better results were obtained using 18 samples for the temporal model. Similarly, in this study the number of samples for EMG time series was low, as it was matched to the number of muscles to allow a fair comparison between the two models. Therefore, the temporal structure can be computed after down-sampling or over-sampling the time series and this may affect the results. The two models exhibit some differences on the datasets employed, probably due to the movements involved. Temporal synergies have been used principally to describe cyclic movements characterized by specific cyclic timings, like locomotion (Ivanenko et al., 2004, 2005). Therefore, in the REACH PLUS dataset, where typical reaching movements are characterized by bell-shaped velocity profile of the end effector trajectory, the temporal synergy model gives results similar to the spatial model. In the NINAPRO dataset, reach to grasp movements are characterized by a triphasic modulation (pre-shaping, grasping and releasing) and more synergies are needed to obtain high *R*^*2*^ reconstruction. Similar results were found also in kinematic synergies applied to hand grasps. Indeed, Jarque-Bou et al. (2019) found that three kinematic synergies accounted for more than the 50% of variance but a high number of synergies was needed to reconstruct finer movements of the hand. Moreover, finer movements of the hand require a more fractionated control of muscles that might be reflected by multiple temporal synergies (Takei et al., 2017).

The temporal synergy structure was analyzed in detail comparing original and surrogate datasets. The differences found in the distribution of the temporal synergies extracted from original and surrogate datasets suggest that the shape of temporal synergies is not simply related to the smoothness of the input signal but it may represent a specific feature of the neural commands. Temporal synergies of the original datasets were narrower and activated closer in time and condensed in the central phase with respect to the smoothed Gaussians of the surrogate data. Thus, neural commands may be generated as a sequence of narrow pulses generated at regular and short intervals during a movement, each activating one or more spatial synergies. The reduction of the full width at half maximum and of the periods when comparing temporal components with the natural composition of a signal with the same frequency content seems to suggest that the configuration of the temporal synergies may reflect an intermittent control of movement. Although the majority of the optimal control theories are based on continuous control signals that generate the human movement, some studies already suggested that the control signal is based on an intermittent control mechanism (Gawthrop et al., 2011; Karniel, 2013). In this control paradigm, the sensory feedback is used intermittently to parameterize the controlled motion law. The CNS sends pulsed commands to generate the movement that are transformed into activation profiles of muscles (Leib et al., 2020) and allows to shape the motor output by adjusting the timing and the amplitude of the bursts (Gross et al., 2002). While the movements analyzed in this study reflect mainly feedforward control (especially in the REACH PLUS dataset), in this study we still observed an intermittent organization of temporal synergies. Given the importance of high dexterity of flexibility of human upper limb and hand, we suppose that the control architecture might be tuned to be intermittent by nature in order to be ready to implement intermittent adaptation and corrections to the movement. In a sensory feedback based experimental design, we expect that intermittent control emerges even further. This is also suggested by the fact that in the NINAPRO (featuring slower movements and higher involvement of the sensory feedback) the difference between the real dataset and the surrogate is amplified with respect to REACH PLUS in which movements are mainly feedforward. This implementation of control of human movement may be related to higher order derivative of the trajectory than the acceleration. Indeed, the intermittent control is predicted by the control signal of the minimum acceleration criterion with constraints (Ben-Itzhak & Karniel, 2008), that is based on minimizing the acceleration while constraining the maximum value of the jerk (third derivative). Intermittent control allows an online optimization process that provides higher adaptability and flexibility of the movement (Loram et al., 2014; van de Kamp et al., 2013).

This finding paves the way for considering spatial and temporal synergies as representative of different features of the modular and hierarchical neuromotor organization. The rationale behind spatial synergies has been widely discussed in the literature. It is based on the observation of spinal modules that produce movements by linear combination of their force or and associated muscle activity output (Bizzi et al., 2002; D’Avella & Bizzi, 2005). This means that spatial modules need divergent neural connection for being implemented and they have been widely related to neural circuits at the spinal level (Saltiel et al., 2001; Tresch et al., 1999). At the same time, in this study we noted that temporal synergies for low orders can achieve higher reconstruction *R*^*2*^. We thus propose that temporal synergies may reflect a neural strategy for generating motor commands, possibly at cortical level, as a reduced number of descending signals recruiting spatial synergies, possibly at spinal level, although Ivanenko et al. (2006) suggested that the temporal patterns of muscle activation during locomotion may also be located in the spinal circuitry. Many studies suggest that CNS structures above spinal level contribute to movement planning. Hart and Giszter (Hart & Giszter, 2004, 2010) demonstrated that the brainstem can be involved in the time-scale distribution, improving smoothness and reducing co-contraction, thus contributing to the implementation of temporal synergies. It has been also demonstrated that internal models controlling the arm movement are located in cerebellum (Kawato, 1999) and multiple inverse and forward models can be adapted to a large set of situations (Haruno et al., 1999; Wolpert & Kawato, 1998). Santello et al. (2013) hypothesized that the cerebellum receives direct input from the spinal premotor pools (and synergies) employed in the tasks. Berger et al. (2020) showed that temporal and spatiotemporal but not spatial structure in the muscle patterns is affected by cerebellar damage. At the same time, the temporal structure of the EMG signals itself can be the result of a set of sensorimotor feedback signals involved in motor control for adaption and tuning (Lockhart & Ting, 2007; Welch & Ting, 2008), thus temporal synergies can be shared between higher and lower levels of the CNS. All these evidences support the notion that synergies are found not only at spinal level. They also contribute to explain the observation that temporal synergies show lower dimensionality with respect to spatial synergies: motion planning takes place in a “principal components-based, synthetic space” that summarizes the main features of the movement more synthetically than the actuation (spatial synergies). Thus, spatial and temporal synergies may not just be “dual models” but they may reflect different levels of a hierarchical organization. Temporal synergies reflect coordination in time, and mapping of goals into high-level features of motor commands (planning); spatial synergies reflect the organization to execute the movement (actuation). Since both models are supported by the data and describe complementary aspects of motor control, a more complete analysis of motor control may be provided by the space-by-time model (Delis et al., 2014, 2015) which incorporates both spatial and temporal synergies. This method identifies both spatial and temporal invariant modules that encode complementary aspects of the tasks: spatial modules identify muscles that activate synchronously, while temporal modules encoded movement phases (Hilt et al., 2018). Spatial synergies can replicate better the original signal, but the temporal model is more efficient at discriminating task (Delis et al., 2018). Overduin et al. (2015) demonstrated that the motor cortex employs spatiotemporal synergies to control movement, indicating that the hierarchical organization of motor control utilizes spatial and temporal features to organize motor synergies. The space-by-time model incorporates both models in compact way, even if it requires more parameters to be stored (Delis et al., 2014). The systematic analysis of spatiotemporal synergies could give a more comprehensive perspective; thus, it should be thoroughly studied in future work.

Understanding better how muscle synergies are related to the neural organization of motor control can be important for many applications. A principal field of application is neurorehabilitation (Singh et al., 2018), since many studies have demonstrated that muscle synergies are a physiological marker in stroke patients (Cheung et al., 2009; Roh et al., 2015). In this way, the rehabilitation can be focused on the altered synergies, improving motor recovery. Furthermore, modularity of motor control can simplify the control of neuroprosthesis (Cole & Ajiboye, 2019; Piazza et al., 2012), allowing to reproduce the desired movement with a small number of command inputs.

## 5. Conclusions

In this paper, we provided an assessment of spatial and temporal synergies and tested their difference in reconstruction accuracy and inter-subject matching. We also showed the existence of both spatial and temporal structure of EMG signal, comparing synergies extracted from the original and a surrogate dataset. While the spatial model has been employed in the vast majority of muscle synergy studies, our results show how the poorly exploited temporal model might also be helpful in the study of motor control. With a detailed characterization of temporal synergies, we suggested that the temporal synergies may capture a higher level of motor organization based on intermittent control that provides flexibility and adaptability of the movement. We believe that this paper will be useful to improve analysis targeting several fields such as rehabilitation, prosthesis control and motor control studies.

## Acknowledgement

Authors wish to thank Robert Mihai Mira for the support in the preliminary part of the experiment.

